# Reactivating Hippocampal-Mediated Memories to Disrupt the Reconsolidation of Fear

**DOI:** 10.1101/2021.09.16.460695

**Authors:** Stephanie L. Grella, Amanda H. Fortin, John H. Bladon, Leanna F. Reynolds, Evan Ruesch, Abby Gross, Monika Shpokayte, Christine Cincotta, Yosif Zaki, Steve Ramirez

## Abstract

Memories are stored in the brain as cellular ensembles activated during learning and reactivated during retrieval. Using the Tet-tag system, we labeled dorsal dentate gyrus (dDG) neurons activated by positive, neutral or negative experiences with channelrhodopsin-2. Following fear-conditioning, these cells were artificially reactivated during fear memory recall. Optical stimulation of a competing positive memory was sufficient to disrupt reconsolidation, thereby reducing conditioned fear acutely and enduringly. Moreover, mice demonstrated operant responding for reactivation of a positive memory, confirming its rewarding properties. These results show that interference from a rewarding experience can counteract negative affective states. While interference induced by memory reactivation involved a relatively small set of neurons, we also found that activating a large population of randomly labeled dDG neurons was effective at disrupting reconsolidation. Importantly, reconsolidation-interference was specific to the fear memory. These findings implicate the dDG as a potential therapeutic node for modulating memories to suppress fear.

## Introduction

Maladaptive conditioned fear, caused by dysregulated fear circuits, plays a significant role in the etiology of anxiety disorders such as post-traumatic stress disorder (PTSD). PTSD can develop in individuals who have experienced a traumatic event and it is often characterized by persistent memories of the trauma^1^. Consequently, contextual fear conditioning (CFC) paradigms have been used as a representative model in animals to study PTSD given it is highly conserved across species^2^. The most widely used CFC paradigms involve pairing an emotionally neutral conditioned stimulus (CS) such as a training context, with an aversive unconditioned stimulus (US) like a foot shock that typically elicits a freezing response in rodents. A learned association emerges, and the CS acquires aversive properties that facilitate retrieval of the conditioned fear memory in the absence of the US. In rodent models, this results in a conditioned fear response upon re-exposure to the context demonstrating this learned relationship^3^. In humans, pathological conditioned fear can occur for decades even in the absence of the exact context in which the traumatic event took place.

In spite of the fact that anxiety disorders are extremely prevalent in the general population, and many individuals experience pathological anxiety as a form of an exaggerated fear state, there are few ways to attenuate maladaptive conditioned fear. Reconsolidation, however, has potential as a therapeutic mechanism for diminishing Pavlovian fear^4^. Reconsolidation theory posits that memories become destabilized during recall as they enter a transient state of malleability where they can be modulated during the time it takes them to restabilize^4, 5^. Despite the long history of experimental reconsolidation-related interventions using a variety of pharmacological agents, behavioral treatments and stimulation protocols to disrupt or enhance memory^6–8^, these studies have yielded mixed results. Only recently has the potential for developing improved reconsolidation-based treatments and novel interventions been recognized ^9, 10^. Nevertheless, most effective therapies for PTSD are trauma-focused, meaning the treatment focuses on the memory of the traumatic event^11^.

Memory is thought to be stored in the sparse activity patterns of neuronal populations within a distributed network^12–14^, or as Wilder Penfield described memory as “the writing left behind the brain by conscious experience”^15^. We often refer to these ensembles, active during memory encoding, as memory traces or engrams^16, 17^ and these engrams are reactivated during retrieval^14, 18^. Findings from several studies have shown that specific memories, including fear memories, can be disrupted by inhibition of associated engrams^12, 19–23^. Specifically, the dorsal dentate gyrus (dDG) of the hippocampus is important for encoding contextual memories^24, 25^, and has been implicated in the pathophysiology of a number of anxiety disorders^25, 26^. Here, we propose a novel intervention based on the hypothesis that using optogenetics to artificially reactivate a previously formed, dDG-mediated memory during reconsolidation will permanently alter and disrupt the original fear memory. We used the Tet-tag system to label dDG neurons activated by exposure to positive, neutral or negative experiences with channelrhodopsin-2 (ChR2)^14, 27, 28^. Mice were subsequently fear conditioned and given a fear memory recall test wherein these tagged neurons were optically reactivated. We hypothesized that this reconsolidation-interference manipulation would update the fear memory with attributes from the interfering engram thereby reducing behavioral expression of conditioned fear. Moreover, as we have previously shown that stress-induced behaviors can be rescued by optically reactivating dDG cells previously active during a positive experience^27^ and others have shown that positive emotions counteract a subset of aftereffects of negative emotions^29^, we proposed that this effect would be more pronounced when the interfering engram was associated with a positive experience compared to a neutral or negative experience.

## Results

### Artificial reactivation of hippocampal-mediated memories during fear memory reconsolidation reduces fear acutely and enduringly

We used a viral, activity-dependent, and inducible neuronal tagging strategy in wild-type c57BL/6 mice. (Fig. 1a). Male mice were injected with virus (either ChR2 or eYFP) and implanted with an optic fiber before being taken off-DOX to open a tagging window^14, 30^. They were split into three groups, and each assigned a differentially-valenced behavioral experience (Fig. 1a). All mice were placed into a novel clean cage and either left undisturbed (neutral)^18^, placed with a female (positive)^27, 31^, or placed into a restraint tube with air holes (negative)^27^ and then placed into the cage. Mice were returned to their home cages 1h later back on DOX to close the tagging window. The following day, mice were fear conditioned in context A and 24h later given a recall test in the same context. During this test, in which we assessed conditioned fear (i.e., freezing) as a proxy for retrieval of the associative fear memory, we simultaneously stimulated the tagged dDG ensembles during the first or last half of the session. We hypothesized that reactivating a positive memory during recall would disrupt reconsolidation potentially altering the fear memory ensemble, and result in decreased freezing at subsequent time points, including during a reinstatement test after an immediate shock in context B. Mice first were fear conditioned using a 4-shock protocol^31, 32^ (Fig. 1b) wherein they exhibited freezing in a stepwise manner, increasing with each successive shock presented (Supplementary Fig. 1). Mice were returned to the same context the next day **(Fig.1c)**. When stimulation occurred during the second half of the session, mice in the positive and negative-ChR2 groups demonstrated a real-time reduction in freezing compared to mice in the neutral-ChR2-group and to eYFP-controls respectively. While there was a natural decline in freezing across the session due to the absence of shock, these mice also showed a significantly steeper decline. No group differences were observed during extinction, immediate shock, or at reinstatement (Fig. 1d-f). However, control mice did freeze significantly more than experimental mice at reinstatement compared to immediate shock (Fig. 1g). In contrast, when mice were fear conditioned (Fig. 1h) and stimulation occurred in the first half of the recall session (Fig. 1i), only mice in the positive- ChR2 group showed reduced fear, which occurred specifically in the last half of the session compared to eYFP-controls. Here, mice in the neutral-ChR2 group demonstrated the fastest extinction rate (Fig. 1j), and as expected, there were no group differences during immediate shock (Fig. 1k). During reinstatement (Fig. 1l), both positive and neutral-ChR2 groups demonstrated reduced fear compared to eYFP-controls, while mice in the negative-ChR2 group did not. Again, control mice froze significantly more than experimental mice at reinstatement compared to immediate shock (Fig. 1m), and this was a larger effect. Therefore, since our effects were greater when stimulation was administered in the first half of the session, we adopted this protocol for all subsequent experiments.

**Fig 1.**
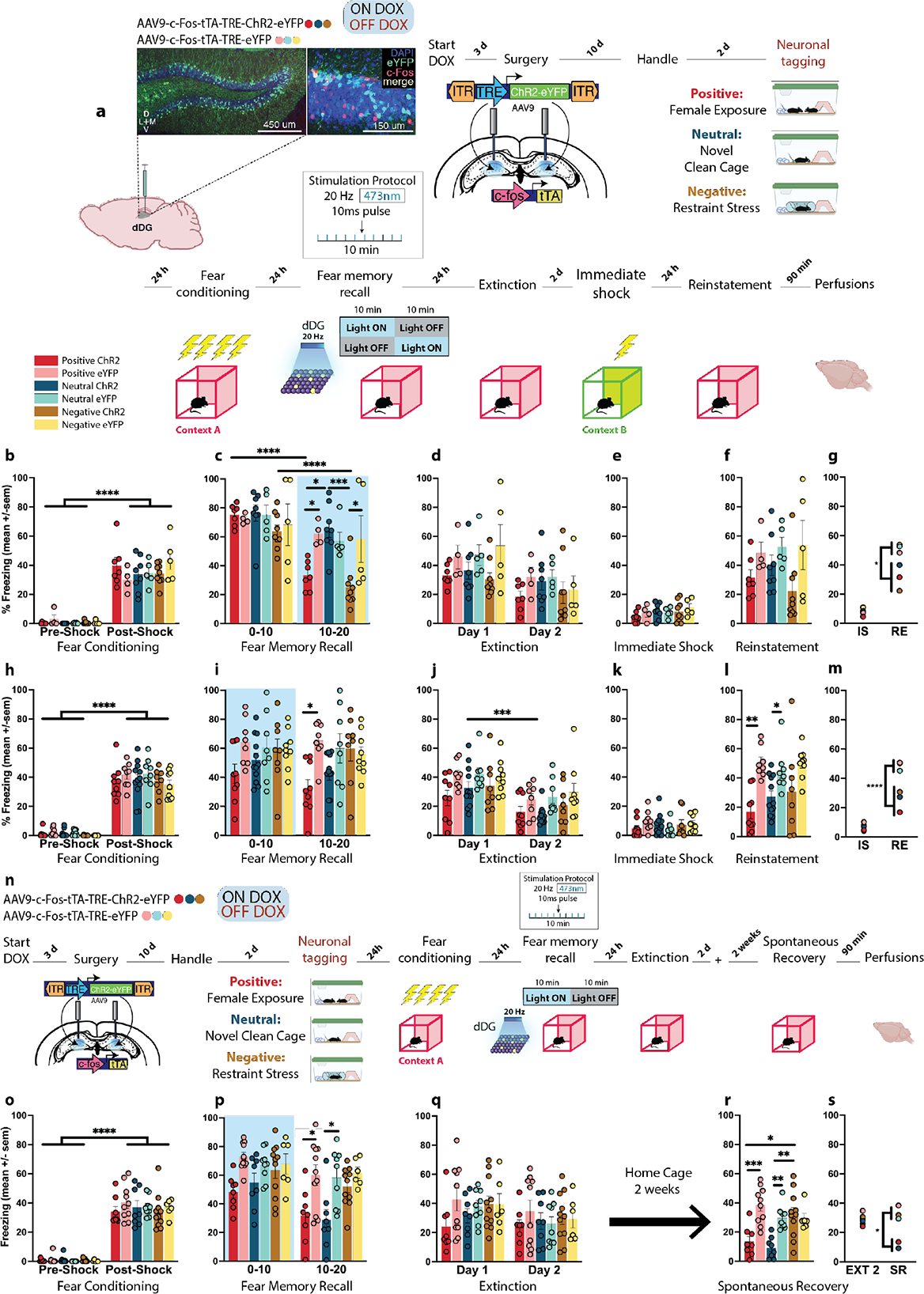
Artificial reactivation of hippocampal-mediated memories during fear memory reconsolidation reduces fear acutely and enduringly. **a,** Schematic of viral strategy and experimental design. dDG cells involved in the encoding of heterogeneously-valenced behavioral epochs (positive, neutral, and negative experiences) were tagged during the off-DOX period (orange). Mice were FC in context A and 24h later given a fear memory recall session during which the cells previously tagged in the dDG were artificially reactivated in either the first, or last half of the session. Across the next two days, mice were given two extinction sessions. The following day, to reinstate fear responding, mice received an immediate shock in context B and were tested for reinstatement the following day. **b,** All mice demonstrated greater freezing post-shock (three-way RM ANOVA: F(1,32)=358.1, P<0.0001, Time). **c,** During recall, when stimulation occurred during the second half of the session, mice in the positive (P=0.0256, P=0.0268) and negative (P=0.0004, P=0.0377) ChR2-groups demonstrated a real-time reduction in freezing compared to mice in the neutral-ChR2 group and to their eYFP-controls respectively (three-way RM ANOVA: F(2,32)=5.143, P=0.0116, Time x Valence x Virus). They also showed a faster decline in freezing across the first and last half of the session (P<0.0001). **d-f,** No group differences during extinction, immediate shock, or when tested at reinstatement. **g,** However, control mice froze significantly more than experimental mice at reinstatement compared to immediate shock (three-way RM ANOVA: F(1,32)=7.362, P=0.0106, Virus x Day). Groups: ChR2 - Positive (n=7 red), Neutral (n=8 turquoise), Negative (n=9 mustard); eYFP - Positive (n=4 pink), Neutral (n=5 cyan), Negative (n=5 yellow). **h,** A separate group of mice were fear conditioned, again demonstrating greater freezing post-shock (three-way RM ANOVA: F(1,48)=761.4, P<0.0001,Time). **i,** When stimulation was given during the first half of the recall session, only mice in the positive-ChR2 group showed reduced fear in the last half of the session compared to eYFP controls (three-way RM ANOVA: F(1,48)=7.737; P=0.0077, Virus). **j,** Mice in the neutral-ChR2 group demonstrated the fastest rate of extinction (three-way RM ANOVA: F(1,48)=57.75, P<0.0001, Time; F(1,48)=10.99; P=0.0017, Virus). **k,** No group differences during immediate shock. **l,** During reinstatement, both positive (P=0.0016) and neutral (P=0.0031) ChR2-groups demonstrated reduced fear compared to eYFP-controls, while mice in the negative-ChR2 group did not (two-way ANOVA: F(1,48)=26.97, P<0.0001, Virus). **m,** Again, control mice froze significantly more than experimental mice at reinstatement compared to immediate shock (three-way RM ANOVA: F(1,48)=24.66, P<0.0001, Day x Virus). Groups: ChR2 - Positive (n=9), Neutral (n=12), Negative (n=8); eYFP - Positive (n=8), Neutral (n=8), Negative (n=9). **n,** In a separate cohort of mice, we assessed the long-term effects of our manipulation. A separate cohort of mice were tested on spontaneous recovery. **o,** Mice were fear conditioned, again demonstrating greater freezing post-shock (three-way RM ANOVA: F(1,46)=685.2, P<0.0001, Time). **p,** Mice in the positive (P=0.0251) and neutral (P=0.0266) ChR2-groups showed reduced fear in the last half of the recall session compared to eYFP controls (three-way RM ANOVA: F(1,46)=16.75, P=0.0002, Time; F(1,46)=28.15, P<0.0001, Virus) **q,** No group differences were observed during extinction. **r,** In a test for spontaneous recovery, mice in both the positive (P=0.0004, P=0.0122) and neutral (P=0.0070, P=0.0011) groups showed reduced freezing compared to their eYFP-controls and compared to the negative-Chr2 group respectively (two-way ANOVA: F(2,46)=6.894, P=0.0024, Valence x Virus). **s,** Positive and neutral ChR2-mice continued to exhibit a reduction in fear two weeks after extinction compared to both positive and neutral-eYFP mice, as well as both groups of negative mice (three-way RM ANOVA: F(2,46)=3.784, P=0.0301, Valence x Virus; F(1,46)=7.859, P=0.0074, Day). Groups: ChR2 - Positive (n=8), Neutral (n=8), Negative (n=11); eYFP - Positive (n=10), Neutral (n=9), Negative (n=6). All data are represented as means ± s.e.m. *P<0.05, **P<0.01, ***P<0.005, ****P<0.00. dDG: dorsal dentate gyrus, DOX: doxycycline, IS: immediate shock, RE: reinstatement.

For the next experiment, we assessed the long-term effects of our manipulation. We replicated the above findings with a similar experimental design. However, instead of giving mice a reinstatement test after immediate shock, we left mice undisturbed in their home cage for 2 weeks after extinction and then gave them a test for spontaneous recovery of fear (Fig. 1n). 24 h after fear conditioning **(Fig.1o)**, during recall, both positive and neutral-ChR2 groups froze less in the last half of the session compared to eYFP-controls (Fig. 1p). All groups extinguished at the same rate (Fig. 1q). Consistent with the effects seen during recall, the test for spontaneous recovery revealed that mice in both the positive and neutral-ChR2 groups froze less compared to eYFP-controls and compared to the negative-Chr2 group demonstrating that our manipulation produced enduring effects on fear memory retrieval processes evident two weeks after extinction (Fig. 1r-s).

### Valence matters: Artificial reactivation of a neutral home cage experience is not sufficient to interfere with fear reconsolidation

The above results illustrate that hippocampal interference resulting from reactivation of positive or neutral engrams is more effective at reducing conditioned fear than engrams associated with a negative experience. To further gauge the importance of valence, first we tested if the novel clean cage experience was indeed “neutral,’’ given that novel stimuli can activate the mesocorticolimbic reward pathway,^33^ and many neurotransmitter systems involved in reward processing^34^. Therefore, for the neutral component of the next experiment, we used a home cage^30^ experience instead where mice were left undisturbed. Secondly, we asked whether a separate positive experience that did not involve male-female interaction was sufficient to reduce fear. Consequently, we used acute cocaine exposure^35^ as our next positive experience. And finally, we asked if the inability to reduce freezing via stimulation of a negative engram during fear memory recall was the result of those memories overlapping. To test this, we tagged dDG cells active when mice were fear conditioned in context C as our next negative experience as this interfering engram would theoretically be composed of some cells of the same cells as the fear memory acquired in context A due to generalization.

To address these questions, we again opened a tagging window off-DOX and labelled a positive, neutral, or negative memory in the dDG (Fig. 2a). Mice assigned to negative groups were initially fear conditioned in context C (Fig. 2b). The following day they were fear conditioned as before in context A (Fig. 2c). Mice fear conditioned the previous day showed higher freezing levels than the other groups pre- and post-shock. During recall (Fig. 2d), mice in the negative groups continued to exhibit more freezing compared to other groups. Additionally, there were no real-time decreases in freezing in any of the ChR2-groups, nor did we see any group differences during the latter half of the recall session. However, mice in positive and neutral-Chr2 groups froze less in comparison to the other groups on extinction day 1 (EXT1) (Fig. 2e). No group differences were observed during immediate shock (Fig. 2f). While we saw a slight decrease in freezing in the neutral-ChR2 group on EXT1, optical stimulation of this home cage memory was not sufficient to compete with the fear memory, as this effect did not persist in our test for reinstatement (Fig. 2g). During reinstatement, we saw that only the positive-Chr2 group had significantly less freezing than eYFP-controls and compared to negative groups. The negative-ChR2 group demonstrated equal freezing levels to controls (Fig 2g-h). These results corroborate our previous findings showing that optical stimulation of a competing positive memory, but not a neutral or negative memory, is sufficient to disrupt reconsolidation of fear.

**Fig. 2.**
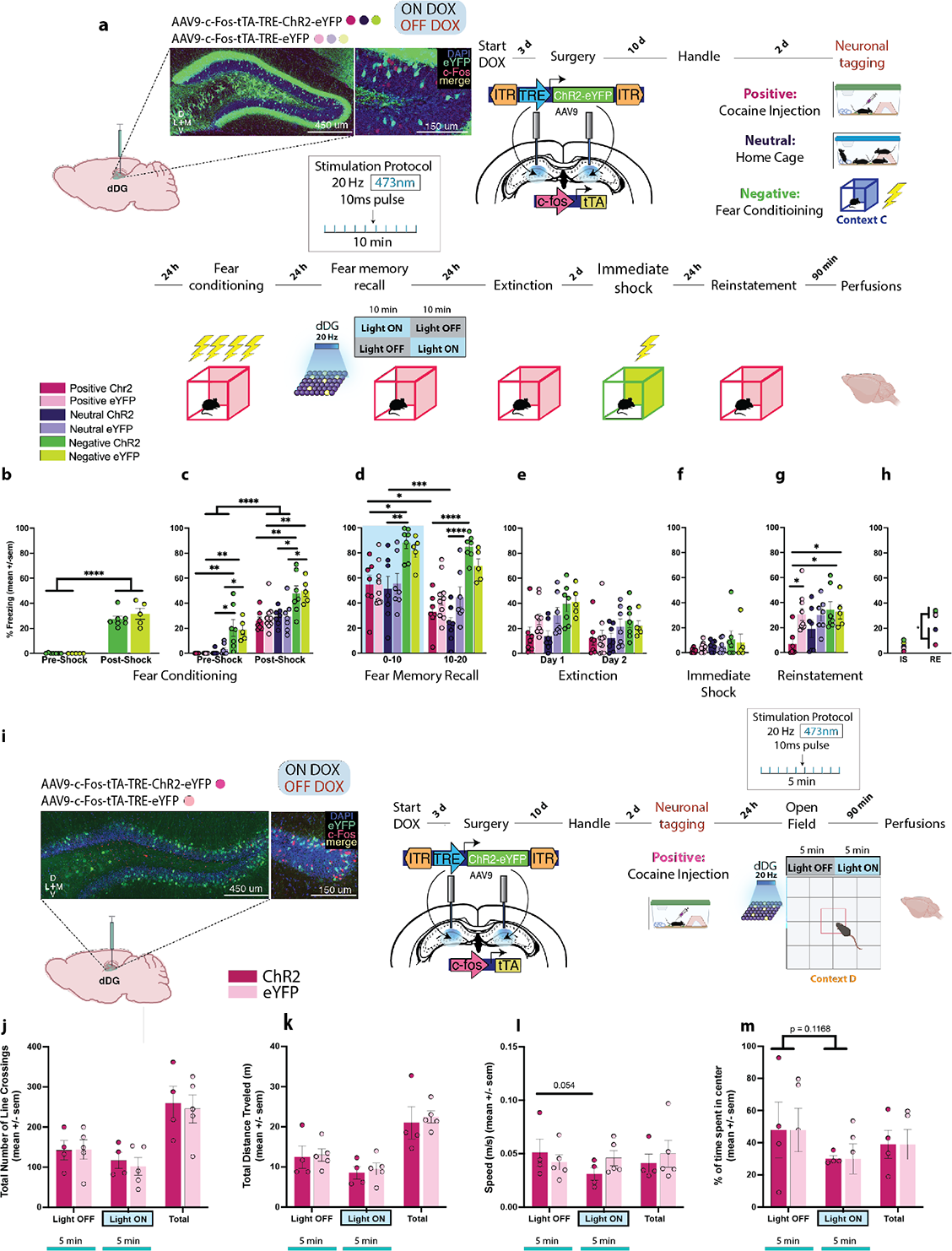
Valence matters: Artificial reactivation of a neutral homecage experience is not sufficient to interfere with fear reconsolidation. **a, Schematic of viral strategy and** experimental design. dDG cells involved in the encoding of heterogeneously-valenced behavioral epochs (positive, neutral, and negative experiences) were tagged during the off-DOX period (orange). The experimental design was the same as before. **b,** Mice in the negative groups were initially fear conditioned in an alternate context (Context F). These mice demonstrated greater freezing post-shock (two-way RM ANOVA: F(1,10)=168.4, P<0.0001, Time). **c,** The following day all mice were fear conditioned and while they all showed increased freezing levels post-shock (three-way RM ANOVA: F(1,37)=295, P<0.0001, Time), mice in the negative groups demonstrated higher freezing levels than the other groups pre-shock (Negative: vs. Positive P<0.0001, vs. Neutral P<0.0001) (two way RM ANOVA: F(2,40)=27.55, P<0.0001, Valence), and post-shock (three-way RM ANOVA: F(2,37)=25.25, P<0.0001, Valence). **d,** During recall, mice in the negative groups continued to exhibit higher levels of freezing compared to the other groups (three-way RM ANOVA: F(2,37)=3.286, P<0.0486, Valence x Virus x Time). Specifically, the negative-ChR2 group froze more than the neutral-ChR2 group in both the first (P=0.0086) and last 10 min (P<0.0001) and also froze more than the positive-Chr2 group at both time points (P=0.255, P<0.0001). Moreover, both the positive (P=0.0072) and neutral (P=0.0002) ChR2 groups showed a faster decline in freezing across time compared to their corresponding eYFP-groups. **e,** During EXT1, mice in the positive and neutral-ChR2 groups froze less than the other groups (three-way ANOVA: F(2,37)=4.107, P=0.0245, Valence x Day; F(2,37)=6.841, P=0.0128, Virus x Day) **f,** No group differences were observed during immediate shock. **g,** During reinstatement, positive ChR2-mice demonstrated reduced fear compared to eYFP-controls (P=0.0249) as well as both the negative-Chr2 (P=0.0147) and eYFP (P=0.0498) groups (two-way ANOVA: F(1,37)=5.923, P=0.0199, Virus; F(2,37)=3.440, P=0.0426, Valence). **h,** Positive and neutral-ChR2 mice froze significantly less than eYFP-mice at reinstatement compared to immediate shock (three-way RM ANOVA: F(1,37)=5.912, P=0.0200, Day x Virus). Groups: ChR2 - Positive (n=7 fuschia), Neutral (n=7 purple), Negative (n=7 green); eYFP - Positive (n=10 pink), Neutral (n=7 violet), Negative (n=5 lime). **i,** Schematic of viral strategy and experimental design. Cells involved in the encoding of an acute, cocaine- exposure (positive) experience were tagged during the off-DOX period (orange). 24h later mice were placed in the OF for 10 min where tagged dDG cells were artificially reactivated in the last 5 min of the session. Measures used to assess locomotion included mean **j,** total number of line crossings **k,** total distance traveled and **l,** speed. No group differences were found suggesting that the reductions in freezing seen in the previous experiments were not due to increased locomotion. **m,** Percentage of time spent in center - no group differences. Groups: ChR2 - (n=4 fuschia), eYFP - (n=5 pink). All data are represented as means ± s.e.m. *P<0.05, **P<0.01, ***P<0.005, ****P<0.00. dDG: dorsal dentate gyrus, DOX: doxycycline, OF: open field, IS: immediate shock, RE: reinstatement.

### The reduction in freezing observed is not due to increases in locomotion

In a separate cohort of mice, dDG cells involved in encoding an acute cocaine-exposure were tagged off-DOX (Fig. 2i). The next day, to assess whether activation of a cocaine engram would induce hyper-locomotor activity, we tested mice in the open field where we reactivated the cocaine engram in the last 5 min of the 10 min test. We found no group differences in total number of line crossings (Fig. 2j), distance traveled (Fig. 2K), or speed **(FIg 2l)** suggesting the decreases in freezing observed in the previous experiment were not due to increased locomotion. Additionally, time spent in the center region revealed no group differences (Fig. 2m) suggesting that artificial activation of a cocaine-related memory is neither anxiogenic, nor anxiolytic.

### Mice will perform an operant response for artificial reactivation of a positive memory

While the hippocampus is implicated in processing positive experiences, it is thought to do so in concert with several regions involved in neuromodulation including the ventral tegmental area (VTA). The VTA is a critical component of the brain’s reward system and negative affective states (e.g., anxiety) are mediated by VTA dysregulation. It is well established that intracranial self-stimulation (ICSS) of the VTA is a powerfully rewarding operant behavior, where rodents maintain delivery of electrical impulses resulting in dopamine release^36, 37^. This procedure has been previously adapted^38–43^ to incorporate *in vivo* optogenetic stimulation of dopaminergic neurons in the VTA. To tag and reactivate dDG cells active during this positive experience, we selectively expressed ChR2 in dopaminergic VTA cells using transgenic mice which express Cre under control of the dopamine transporter (DAT). We injected our viral vectors AAV5-Ef1a-DIO-(hChR2-E123A)-eYFP and implanted an optic fiber unilaterally into the VTA. We also injected c-Fos-tTA-TRE-(ChR2)-eYFP and implanted optic fibers bilaterally aimed at the dDG (Fig. 3a). Mice were initially habituated to the operant chamber and given access to a wheel with no consequences, which served as a baseline measure. The following day, mice were placed back into the operant box and two nose ports were introduced, one active and one inactive. Nose pokes into the active port produced optogenetic VTA stimulation while nose pokes into the inactive port produced no stimulation and served as a discriminative control. Mice were given three ICSS training sessions and then taken off-DOX. They were brought back for a fourth training session, in which dDG cells responsive to VTA self-stimulation were tagged. The following day, access to the nose ports was restricted and spins on the wheel produced optical stimulation of the dDG to reactivate the VTA self-stimulation engram. This was done to assess whether mice would perform an operant response for a positive (VTA-ChR2, dDG-ChR2) or neutral (VTA-eYFP, dDG-ChR2) experience compared to dDG-eYFP controls (Fig. 3a). In mice injected with ChR2 in the VTA, nose pokes into the active port were significantly higher than the inactive port, and they increased across sessions demonstrating the mice’s ability to discriminate between ports and self-deliver optical stimulation for reward (Fig. 3b-e). Comparing wheel baseline measures to training and test, mice injected with ChR2 in the VTA and dDG, completed significantly more wheel rotations, which were kept in motion for longer durations and distances (Fig. 3f-h) and produced more stimulations (Fig. 3i) compared to all other groups. This finding demonstrates that mice will perform an operant response to maintain artificial reactivation of a positive memory, specifically the memory of VTA self-stimulation. Mice did not exhibit this behavior for a memory of operant box exposure in the absence of VTA stimulation. Following this test, mice underwent the same experimental protocol as before where they were fear conditioned in context A (Fig. 3j) and then given a recall test. Reactivation of the VTA self-stimulation engram (VCDC) during recall significantly reduced freezing levels throughout the session (Fig. 3k). Levels remained low throughout extinction (Fig. 3l), immediate shock (Fig. 3m), and reinstatement (Fig. 3n). Interestingly, between the two dDG-eYFP groups, the group that had received VTA stimulation earlier (VCDE) demonstrated a beneficial effect of this experience exhibiting intermediate levels of freezing compared to the VCDC group and the other control groups on EXT1 (Fig 3l), and from immediate shock to reinstatement (Fig. 3o). Together, these results show that interference from a rewarding experience can counteract negative affective states.

**Fig. 3.**
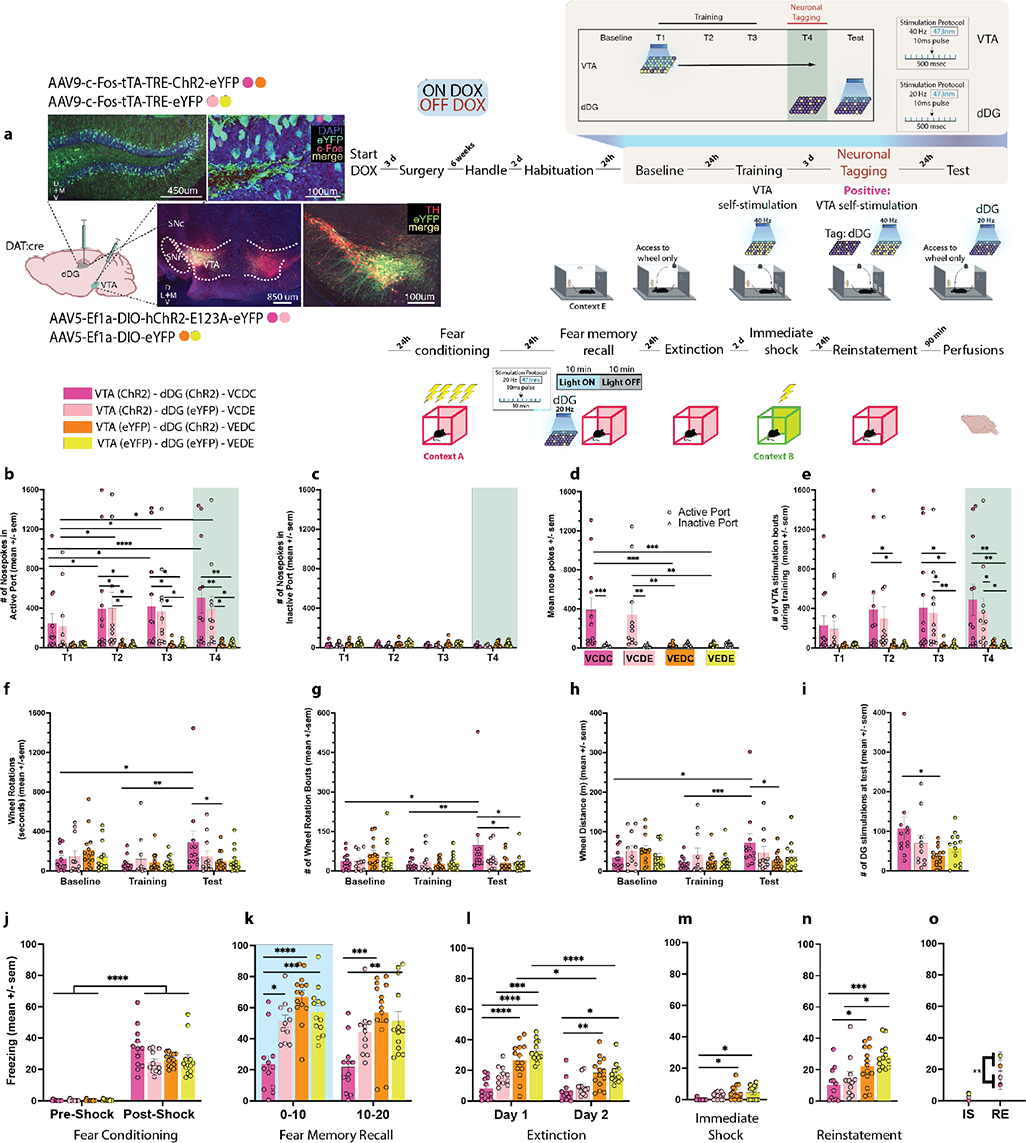
Mice will perform an operant response for artificial reactivation of a positive memory. **a,** Schematic of viral strategy and experimental design. DAT Cre mice were initially given access to a wheel that produced no consequences when it was spun (baseline). They were then trained to nose poke for closed-loop optogenetic self-stimulation of the VTA for 500 ms bursts of stimulation (T1-T3). dDg cells involved in the encoding of this positive experience were tagged during the off-DOX period (orange, T4). The following day, mice were again given access to the wheel, where spinning it produced artificial reactivation of the tagged dDG cells responsive to VTA self-stimulation. The next day, mice underwent the same experimental procedure as the previous experiments. Groups: Mice were injected with either ChR2 (VC) or eYFP (VE) in the VTA, and also injected with either ChR2 (DC) or eYFP (DE) in the dDG. Groups: VCDC (n=12 dark pink), VCDE (n=11 light pink), VEDC (n=14 orange), VEDE (n=13 yellow). **b,** In the active port, VCDC & VCDE mice nose-poked significantly more than VEDC & VEDE mice (two-way RM ANOVA: F(9,138)=2.179, P=0.0270, Time x Group). These mice also increased responding across days whereas **c,** no effects were seen in the inactive port. **d,** Summary of nose poke behavior across days for each group (two-way RM ANOVA: F(3,46)=6.102, P=0.0014, Noseport x Group). **e,** Nose pokes into the active port resulted in VTA stimulation bouts (500ms), which were observed significantly more in the VCDC and VCDE groups (two-way RM ANOVA: F(9,138)=2.472, P=0.0120, Time x Group). **f,** During baseline and training wheel spins produced no consequences. During the test session wheel spins resulted in reactivation of dDG cells involved in encoding the last VTA self-stimulation session (T4). We measured the number of wheel rotations and found that mice in the VCDC group spun the wheel for significantly longer than than VEDC mice during the test (P=0.0264) and during baseline (P=0.0230) and training (P=0.0029) demonstrating that they will perform an operant response for access to a positive memory (two-way RM ANOVA: F(6,92)=2.233, P=0.0467, Time x Group). **g,** We found a similar tendency for the number of times the wheel was spun (At test - VCDC vs. VEDC: P=0.0108, VCDC vs. VEDE: P=0.0133; VCDC - from baseline to test: P=0.0148, from training to test: P=0.0032) (two-way RM ANOVA: F(6,92)=2.513, P=0.0269, Time x Group). **h,** This relationship was also observed when looking at wheel distance (At test - VCDC vs. VEDC: P=0.0246; VCDC - from baseline to test: P=0.0177, from training to test: P=0.0009) (two-way RM ANOVA: F(6,92)=2.575, P=0.0237, Time x Group). **i,** Finally, we found that VCDC mice produced significantly more dDG stimulations at test than the VEDC group (P=0.0318) (one-way ANOVA: F(3,46)=2.836, P=0.0484). **j,** The following day all mice were fear conditioned and showed increased freezing post-shock (three way RM ANOVA: F(1,92)=386.4, P<0.0001, Time). **k,** During recall, stimulation produced real-time decreases in freezing in the VCDC group compared to all other groups (vs. VCDE: P=0.0112, VEDC: P<0.0001, VEDE: P=0.0004). This persisted in the last half of the session (10-20) (vs. VEDC: P=0.0001, VEDE: P=0.0027) (three-way RM ANOVA: F(1,46)=11.65, P=0.0013, dDG Virus x VTA Virus; F(1,46)=9.841, P=0.003, Time). **l,** During extinction, VCDC mice showed reduced freezing compared to VEDC (Day 1: P<0.0001, Day 2: P=0.0090) and VEDE (Day 1: P<0.0001, Day 2: P=0.0144) controls (three way RM ANOVA: F(1,46)=8.526, P=0.0054, VTA Virus x Day; F(1,46)=5.896, P=0.0191, dDG Virus x Day). **m,** These same differences were seen during immediate shock (VCDC vs. VEDC: P=0.0107; VCDC vs. VEDE: P=0.0201) (two-way ANOVA: F(1,46)=10.70, P=0.002, VTA Virus). **n,** During reinstatement, VCDC mice demonstrated reduced fear compared to VEDC (P=0.0405) and VEDE controls (P=0.0007). Interestingly, VCDE mice, which differed from VEDE mice by their VTA self-stimulation experience alone, also showed less freezing (P=0.0199) (two-way ANOVA: F(1,46)=16.78, P=0.0002, VTA Virus). **o,** VCDC and VCDE mice froze significantly less than VTA-eYFP controls (VEDC & VEDE mice) at reinstatement compared to immediate shock (three-way RM ANOVA: F(1,46)=7.879, P=0.0073, VTA Virus x Day). All data are represented as means ± s.e.m. *P<0.05, **P<0.01, ***P<0.005, ****P<0.00. dDG: dorsal dentate gyrus, DOX: doxycycline, IS: immediate shock, RE: reinstatement, VTA: ventral tegmental area.

### Activation of randomly labeled dDG neurons is also sufficient to disrupt fear reconsolidation

Next we asked, if reconsolidation-interference induced by reactivation of a positive memory involves a small set of neurons (<10%)^30^, could we circumvent the positive-valence prerequisite to achieve the same effect if we activated a large population of neurons not necessarily tied to a memory. Unlike the previous experiments, where we used a cFos-inducible tagging strategy to label cells involved in different experiences, here, we used a virus with a constitutive promoter (CaMKIIa) to randomly tag dDG neurons with ChR2 (Fig. 4a). Mice were injected with either undiluted or diluted virus to label a large percentage or fraction of dDG cells, respectively. Mice were fear conditioned (Fig. 4b) and during recall the next day, the labeled neurons were optically stimulated. During the first half of the session, we saw real-time decreases in freezing in both undiluted and diluted-ChR2 groups but by the second half of the session only the undiluted group showed significantly less freezing (Fig. 4c). The undiluted- Chr2 group continued to exhibit less freezing on EXT1 (Fig. 4d) and there were no group differences in immediate shock (Fig. 4e). At reinstatement, we observed reduced freezing in both undiluted and diluted-ChR2 groups compared to eYFP-controls (Fig. 4f) and compared to immediate shock (Fig. 4g). Our effects were greater in the undiluted group, suggesting reconsolidation-interference can be effectively achieved by activating ensembles that are not connected to an engram of a particular valence if enough cells are activated. This memory modulation strategy may be akin to stimulation protocols currently approved for use in humans^44, 45^.

**Fig. 4.**
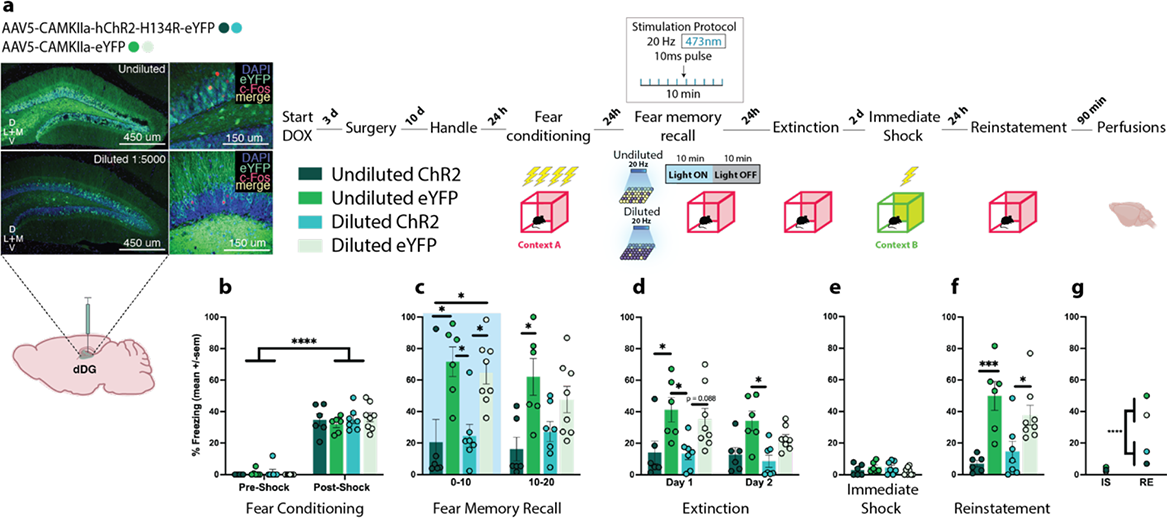
Activation of randomly labeled dDG is also sufficient to disrupt fear reconsolidation. **a,** Schematic of viral strategy and experimental design. We used a virus with a constitutive promoter to randomly label dDG neurons not tied to the encoding of a behavioral epoch. Mice were injected with either undiluted or diluted virus. The experimental procedure was similar to the previous experiments with the exception of the neuronal tagging component. Groups: Undiluted - ChR2 (n=6 teal), eYFP (n=6 green); Diluted - ChR2 (n=7 aqua), eYFP (n=8 mint green) **b,** During fear conditioning, all mice froze post-shock (three-way RM ANOVA: F(1,23)=447.9, P<0.0001, Time)**. c,** During recall, stimulation produced real-time decreases in freezing in both the undiluted (P=0.0106) and diluted (P=0.0369) ChR2-groups compared to eYFP-controls (three-way RM ANOVA: F(1,23)=25.67, P<0.0001, Virus). In the latter half of the session, mice in the undiluted ChR2 continued to show decreased freezing compared to eYFP- controls (P=0.0300). **d,** During extinction, there was an overall decrease in freezing across days (three-way RM ANOVA: F(1,23)=4.312, P=0.0492, Time) and a reduction in freezing in ChR2- groups compared to eYFP-controls (three-way ANOVA: F(1,23)=20.88, P=0.0001, Virus). More specifically, in the undiluted-groups on day 1 (P=0.0394). **e,** No group differences during immediate shock. **f,** During reinstatement, mice in both the undiluted (P=0.0004) and diluted (P=0.0304) ChR2-groups froze less compared to eYFP-controls (two-way ANOVA: F(1,23)=25.15, P<0.0001, Virus). **g,** They also froze significantly less at reinstatement compared to immediate shock (three-way RM ANOVA: F(1,23)=27.08, P<0.0001, Virus x Day). All data are represented as means ± s.e.m. *P<0.05, **P<0.01, ***P<0.005, ****P<0.00. dDG: dorsal dentate gyrus, IS: immediate shock, RE: reinstatement.

### dDG interference is specific to the fear memory and does not affect other hippocampal- mediated memories

To determine whether our manipulation, which disrupts the fear memory, would also indiscriminately affect other types of hippocampal-mediated memory, we trained mice on a spatial reference memory task. Like the previous experiment, we injected mice with undiluted AAV5-CaMKIIa-(hCHR2-H134R)-eYFP to randomly label dDG neurons (Fig. 5a). Water-restricted mice were then trained to obtain a water reward from one of the arms in an 8-arm radial maze. They received 4 trials / d until a set of performance criteria were met, which took between 5-14 days (Fig. 5b). Afterwards, they underwent the same behavioral protocol as the previous experiments. We retested mice on spatial memory performance after fear conditioning and after recall. Following fear conditioning (Fig. 5c), during recall, ChR2-mice showed real-time decreases in freezing (Fig. 5d) which persisted in extinction (Fig. 5e). No differences were seen during immediate shock (Fig. 5f), and ChR2-groups demonstrated less freezing during reinstatement (Fig. 5g) and from immediate shock to reinstatement (Fig. 5h). For maze performance, all mice improved across time in terms of latency to find the reward, number of arm-deviations (upon a mouse’s first visit to an arm), number of reference errors (i.e., entering the wrong arm) and repeated reference errors (working memory errors) made (Fig. 5i-p **left panels)**. Performance was divided into two categories: trial 1, which was interpreted as an assessment of long-term memory from the day before, and trials 2-4, which were interpreted as an assessment of short-term memory from the previous trial on the same day. Mice were first tested in the maze 3h after fear conditioning to confirm that fear conditioning itself did not affect spatial memory. We saw no effects (Fig. 5i-p **middle panels)**.

**Fig. 5.**
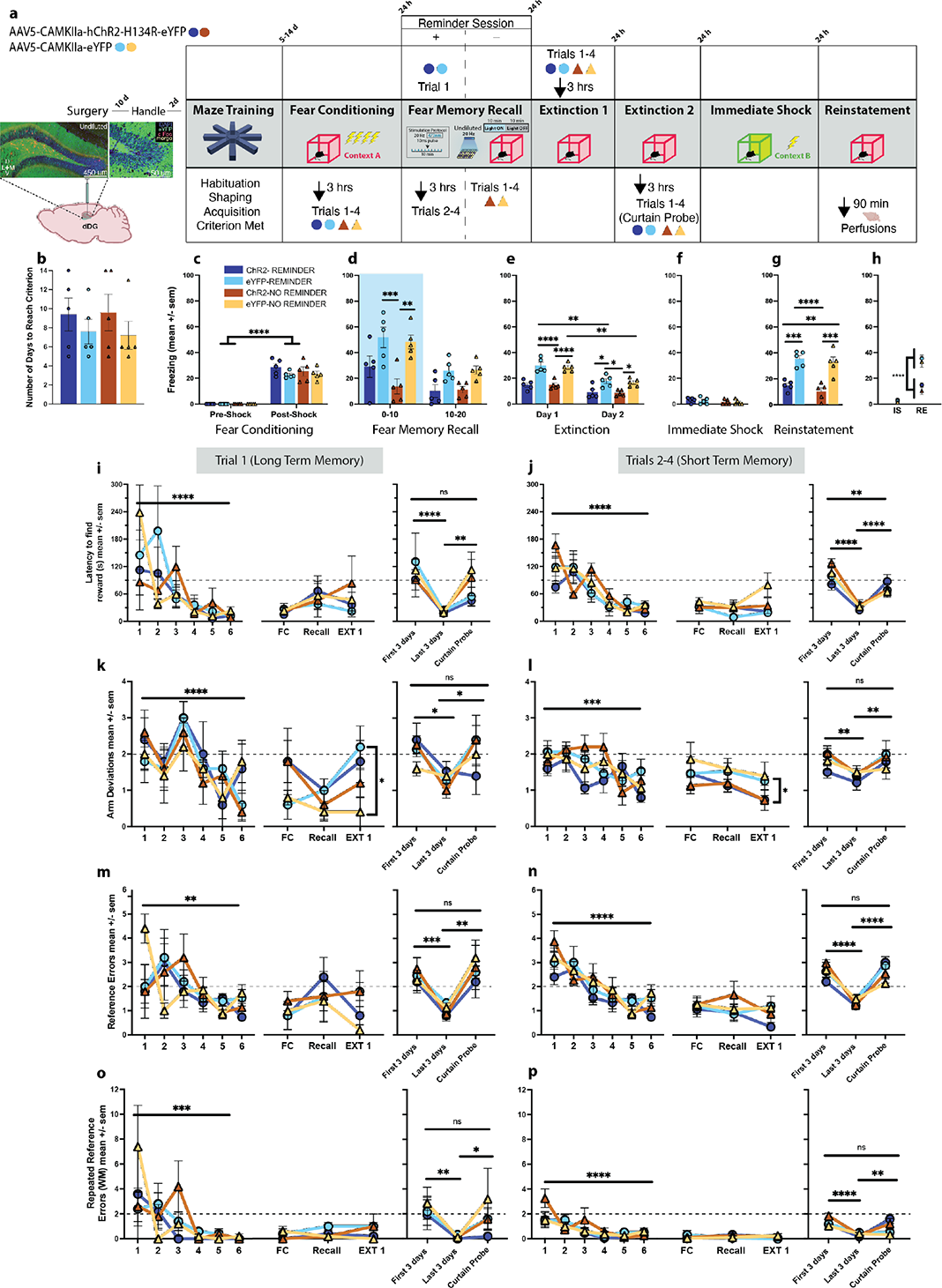
dDG interference is specific to the fear memory and does not affect other hippocampal-mediated memories. **a,** Schematic of viral strategy and experimental design. To determine whether our manipulation affects other types of hippocampal-mediated memory, we trained mice on a spatial reference task. Mice were injected with undiluted AAV5-CaMKIIa- (hCHR2-H134R)-eYFP virus to randomly label dDG neurons. Water-restricted mice were trained to obtain a water reward from one of the arms in an 8-arm radial maze (4 trials / day until criterion). Once criterion was met, they were FC as before and 3h later performance on the maze was tested to assess whether FC affected spatial memory. The following day they received a fear memory recall test, and as before, stimulation of the labeled neurons. Half the mice received (Reminder groups) one reminder session (trial 1) prior to recall and trials 2-4 3h after the recall session. The other half of mice (No Reminder groups) did not receive a reminder session, and instead received all 4 regular trials 3h after the recall session. All mice were tested on the maze the following day and 3h later were given the first EXT session. The following day mice received the second EXT session and 3 h later were given a curtain probe test. The next day, mice underwent immediate shock in context B and 24 h later a reinstatement test. Groups: Reminder - ChR2 (n=5 dark blue), eYFP (n=5 light blue); No Reminder - ChR2 (n=5 bronze), eYFP (n=5 gold). **b,** Mice took between 5-14 days to reach criterion. **c,** During FC, all mice froze post-shock (three-way RM ANOVA: F(1,16)=377.1, P<0.0001, Time)**. d,** During recall, stimulation produced real-time decreases in freezing in the in the No Reminder-ChR2 group (P=0.0027) compared to the eYFP control (three-way RM ANOVA: F(1,16)=26.11, P=0.0001, Time; F(1,16)=24.75, P=0.0001, Virus)**. e,** During EXT, there was an overall decrease in freezing across days and a reduction in freezing in ChR2-groups compared to eYFP-controls (three-way RM ANOVA: F(1,16)=6.008, P=0.0261, Time x Virus). More specifically, in the No Reminder group on day 1 (P<0.0001) and in both the Reminder (P=0.0279) and No Reminder (P=0.0395) groups on day 2. **f,** No group differences during immediate shock. **g,** During reinstatement, mice in both the Reminder (P=0.0008) and No Reminder (P=0.0003) ChR2 groups froze less compared to eYFP controls (two-way ANOVA: F(1,16)=53.27, P<0.0001, Virus). **h,** They also froze significantly less at reinstatement compared to immediate shock (three-way RM ANOVA: F(1,16)=66.81, P<0.0001, Virus x Day). **i,** For Trial 1 (LTM): Latency to find reward. Left panel - first 3 and last 3 days of training; All mice completed the task more quickly across time (three-way RM ANOVA: F(5,48)=8.053, P<0.0001, Day). Middle panel - No differences across sessions on FC, Recall, and EXT 1 days. Right panel - Mice performed just as poorly during the curtain probe as they did when they initially began training (F3 vs L3: P<0.0001; L3 vs CP: P=0.0041) (three-way RM ANOVA: F(2,24)=15.16, P<0.0001, Time). **j,** For Trials 2-4 (STM): Latency to find reward. Left panel - All mice completed the task more quickly across time (three-way RM ANOVA: F(5,48)=19.66, P<0.0001, Day). Middle panel - No differences across sessions. Right panel - Mice performed just as poorly during the curtain probe as they did when they initially started training (F3 vs L3: P<0.0001; L3 vs CP: P<0.0001) (three-way RM ANOVA: F(2,24)=43.93, P<0.0001, Time). **k,** Arm Deviations - Trial 1: Left - All mice completed the task more quickly across time (three-way RM ANOVA: F(5,48)=6.817, P<0.0001, Day). Middle - Both ChR2 and eYFP-mice in the Reminder groups had higher arm deviations during the first trial prior to the first fear memory EXT session (three-way RM ANOVA: F(1,24)=4.741, P=0.0395, Reminder). The Reminder session briefly affected this measure of spatial memory. This was not a result of our reconsolidation interference protocol as there were no differences between ChR2 and eYFP-mice. Right - curtain probe diminished performance (F3 vs L3: P=0.0162; L3 vs CP: P=0.0249) (three-way RM ANOVA: F(2,24)=5.671, P=0.0096, Time). **l,** Arm Deviations - Trials 2-4: Left - All mice completed the task more quickly across time (three-way RM ANOVA: F(5,48)=5.186, P=0.0007, Day). Middle - Mice in the ChR2- groups (both Reminder and No Reminder) demonstrated better performance (fewer arm deviations) compared to eYFP-groups on EXT 1 suggesting that our manipulation improved performance on the maze despite whether a reminder session was given (three-way RM ANOVA: F(1,24)=6.819, P=0.0153, Virus). Right - curtain probe diminished performance (F3 vs L3: P=0.0036; L3 vs CP: P=0.0045) (three-way RM ANOVA: F(2,24)=8.610, P=0.0015, Time). **m,** Reference Errors - Trial 1: Left - All mice completed the task more quickly across time (three-way RM ANOVA: F(5,48)=3.921, P=0.0046, Day). Middle - No differences across sessions. Right - curtain probe diminished performance (F3 vs L3: P=0.0012; L3 vs CP: P=0.0002) (three-way RM ANOVA: F(2,24)=13.80, P=0.0001, Time). **n,** Reference Errors - Trials 2-4: Left - All mice completed the task more quickly across time (three-way RM ANOVA: F(5,48)=14.94, P<0.0001, Day). Middle - No differences across sessions. Right - curtain probe diminished performance (F3 vs L3: P<0.0001; L3 vs CP: P<0.0001) (three-way RM ANOVA: F(2,24)=6.009, P<0.0077, Time x Reminder). **o,** Repeated Reference Errors - Trial 1: Left - All mice completed the task more quickly across time (three-way RM ANOVA: F(5,48)=6.392, P=0.0001, Day). Middle - No differences across session. Right - curtain probe diminished performance (F3 vs L3: P=0.0018; L3 vs CP: P=0.0455) (three-way RM ANOVA: F(2,24)=7.927, P=0.0023, Time). **p,** Repeated Reference Errors - Trials 2-4: Left - All mice completed the task more quickly across time (three-way RM ANOVA: F(5,48)=9.848, P<0.0001, Day). Middle - No differences across sessions. Right - curtain probe diminished performance (F3 vs L3: P<0.0001; L3 vs CP: P=0.0083) (three-way RM ANOVA: F(2,24)=5.862, P=0.0085, Time x Virus). Dotted lines represent the criterion used; mice had to meet this in each measure for 2 consecutive days to reach performance criterion. All data are represented as means ± s.e.m. *P<0.05, **P<0.01, ***P<0.005, ****P<0.00. dDG: dorsal dentate gyrus, EXT: extinction, FC: fear conditioning, HPC: hippocampus, IS: immediate shock, LTM: long-term memory, STM: short term memory, RE: reinstatement.

Working under the assumption that the disruption of the fear memory by our interference manipulation occurs because the conditions are present for memory to undergo reconsolidation, observing an impairment of spatial memory could be obscured if this memory does not also undergo reconsolidation. To accurately test whether our interference manipulation is specific to the fear memory, we sought to increase the likelihood that boundary conditions permitting memory reconsolidation for the spatial memory were met^46^. To that end, we gave half the mice (Reminder groups) one reminder session (trial 1) prior to the fear memory recall test. During this reminder session, we replaced the flooring in the maze center with sandpaper to create a prediction error^47^. This did not affect reward-location, but it did present mice with an unexpected cue during the trial. Reminder mice received trials 2-4 3h after the recall session while the other half of mice (No Reminder groups) did not receive a reminder session, and instead received all 4 regular trials 3h after the recall session. Again, we saw no differences in performance and all mice performed well within the range of criterion (Fig. 5i-p **middle panels)**. All mice were tested on the maze the following day and 3h later were given the first extinction session. No measures were affected during this test except arm-deviations. On trial 1, both ChR2 and eYFP mice in the Reminder groups had higher arm-deviations suggesting the reminder session, which potentially led to reconsolidation, briefly affected this measure of spatial memory. This effect was behavioral and not a result of reconsolidation-interference as there were no differences between ChR2 and eYFP-mice. On trials 2-4, mice in ChR2-groups (both Reminder and No Reminder) demonstrated better performance (fewer arm-deviations) compared to eYFP-groups suggesting reconsolidation-interference actually improved maze performance regardless of whether a reminder session was given. Importantly, this also demonstrated that the disruptive effect of our manipulation was specific to the fear memory (Fig. 5i-p **middle panels)**.

To ensure we were testing hippocampal-mediated memory where mice were using extra-maze cues to find the reward, the following day mice were given a curtain probe test. All mice showed decreased performance on the probe compared to the last 3 d of training with performance reaching the level it was at when they first started the task (Fig. 5i-p **right panels).**

### Rewriting the original fear memory

Next, we sought to determine whether the reduction in conditioned fear was accompanied by a change in ensemble dynamics of the original fear engram. To assess whether our manipulation had altered the original fear memory we combined two virus-based systems (Fig. 6a). All mice were injected with c-Fos-tTA-TRE-mCherry to tag dDG cells involved in encoding the fear conditioning epoch. Mice were also injected with either undiluted AAV5-CaMKIIa-hCHR2-H134R-eYFP or undiluted AAV5-CaMKIIa-eYFP to label an “interfering” set of dDG neurons. Mice then underwent the same experimental protocol as before (Fig. 6b). During recall, stimulation produced real time decreases in freezing in ChR2-mice compared to eYFP-controls (Fig. 6c) and these differences persisted in extinction (Fig. 6d). No group differences were observed during immediate shock (Fig 6e). During reinstatement, mice in the ChR2-group continued to show decreased levels of freezing (Fig. 6f) and froze significantly less at reinstatement compared to immediate shock (Fig. 6g). To determine whether these mice had fewer overlapping neurons from the original fear conditioning epoch to the reinstatement test compared to eYFP-controls, mice were perfused 90 min after reinstatement and c-Fos levels were quantified.

**Fig. 6.**
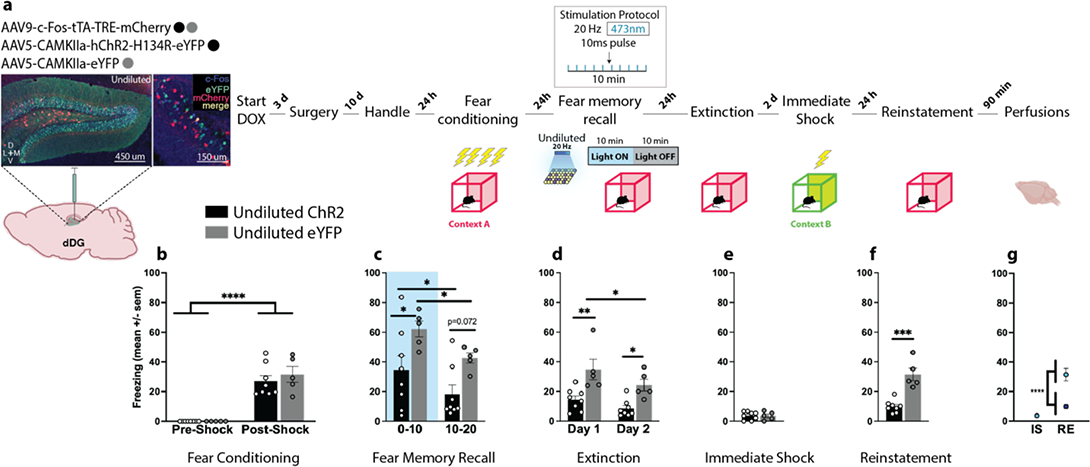
A tale of two memories. **a,** Schematic of viral strategy and experimental design. To assess whether our reconsolidation interference manipulation was able to alter the original fear engram, we combined two viral strategies. All mice were injected with c-Fos-tTA-TRE-mCherry to tag dDG cells involved in the encoding of the fear conditioning epoch. Mice were also injected with either undiluted AAV5-CaMKIIa-hCHR2-H134R-eYFP (n=8, black) or undiluted AAV5- CaMKIIa-eYFP (n=5, grey) to randomly label dDG neurons. Mice then underwent the same experimental protocol as before. **b,** All mice froze post-shock (two-way RM ANOVA: F(1,11)=85.26, P<0.0001, Time)**. c,** During recall, stimulation produced real-time decreases in freezing (0-10) (P=0.0392) in ChR2-mice compared to eYFP-controls (two-way RM ANOVA: F(1,11)=20.32, P=0.0009, Time; F(1,11)=6.468, P=0.0273, Virus). **d,** These differences persisted in extinction (Day 1: P=0.0016; Day 2: P=0.0127) (two-way RM ANOVA: F(1,11)=15.05, P=0.0026, Time; F(1,11)=14.26, P=0.0031, Virus). **e,** No group differences were observed during immediate shock. **f,** During reinstatement, mice in the ChR2 group continued to show decreased levels of freezing (paired t-test: t(11)=5.768, P=0.0001, two-tailed). **g,** They also froze significantly less at reinstatement compared to immediate shock (two-way RM ANOVA: F(1,11)=22.40, P=0.0006, Virus x Day). All data are represented as means ± s.e.m. *P<0.05, **P<0.01, ***P<0.005, ****P<0.00. dDG: dorsal dentate gyrus, IS: immediate shock, RE: reinstatement.

For each experiment, representative images from each neuronal tagging strategy were taken (Fig. 7a-d; Supplemental Fig. 2a-e) and used to obtain cell counts. We first calculated the total number of DAPI labeled cells for each group (Fig. 7e-f). For experiments where we tagged a behavioral epoch using a c-Fos promoter (Fig. 1-3, 6), the size of the engram was determined as a percentage of DAPI-labeled neurons (Fig. 7g**, eYFP;** Fig. 7h**, mCherry)**. Regardless of valence, tagged cells associated with a particular behavioral experience, including a home cage experience^30^, consisted of approximately 8% of DAPI-labeled neurons in the dDG (M=8.015, SD=0.572). The percentage of cells labeled as c-Fos^+^, representing the set of cells active during the reinstatement test (Fig. 7g**&i, RFP;** Fig. 7h**&j, RFP & BFP)** was approximately 1.5% (M=1.438, SD=0.21). In the first set of experiments, the percentage of overlap (yellow) (Fig. 7g**&k)** between the tagged engram used to interfere with the fear memory during recall (eYFP) (Fig. 7g) and the engram at reinstatement (RFP) (Fig. 7g**&i**) can be interpreted as a measure of whether our reconsolidation-interference manipulation resulted in merging the two memories together. We observed a greater percentage of overlap for negative experiences compared to neutral experiences suggesting the negative experiences shared higher similarity with the fear memory compared to the neutral experiences. There were no differences between CHr2 and eYFP mice, however, leading us to conclude that stimulation of an interfering engram during recall did not increase the similarity between the engram and the fear memory at reinstatement (Fig. 7k). Similarly, when we tagged neurons not involved in the encoding of a behavioral epoch, we saw a similar trend where overlaps did not differ between ChR2 and eYFP groups suggesting that activation of the randomly labeled neurons did not then become recruited into the fear memory engram at reinstatement (Fig. 7h**&l)**. However, there was a higher degree of overlap in the undiluted groups compared to the diluted groups based simply on the number cells tagged. In the undiluted groups, we tagged approximately 40% (M=37.16, SD=5.5) of DAPI-labeled neurons and in the diluted groups we tagged a similar 8% of cells (M=8.57, SD=0.4) as our epoch-associated engrams (Fig. 7h). Finally, we sought to determine whether the original fear memory was changed from conditioning to reinstatement given the reduction in fear, despite not being biased to incorporate the cells artificially activated during recall. The percentage of overlaps (magenta) (Fig. 7h**&l)** between these sets of neurons, those tagged in the original fear engram (mCherry) (Fig. 7h) and the fear engram at reinstatement (BFP) (Fig. 7h**&j)** can be interpreted as a measure of whether reconsolidation-interference resulted in alteration of the original fear memory. We found significantly fewer overlaps in the fear memory across conditioning and reinstatement in the ChR2-group, suggesting that while we did not merge the fear memory with the interfering ensembles (Fig. 7m), our manipulations, nonetheless, altered the original fear memory (Fig. 7n), which we believe may help explain the corresponding reductions in fear. We speculate that our reconsolidation-interference manipulation caused a disengagement of these ensembles in an orthogonal manner, separating these memories into disparate neuronal populations (Fig. 7m-n). Interestingly, the proportion of randomly labeled / activated (eYFP) cells that were also part of the original fear engram (mCherry) was above chance (yellow) and the proportion of randomly labeled / activated (eYFP) cells that were also part of the fear engram at reinstatement (BFP) was also above chance (cyan) as was overlap of all 3 types of cells (white) in the eYFP group (Fig. 7l); however, no group differences were seen. Taken together, these findings provide preliminary evidence for the potential therapeutic efficacy of artificially modulating memories to both acutely and enduringly suppress fear responses by altering the original fear memory during the reconsolidation period.

**Fig. 7.**
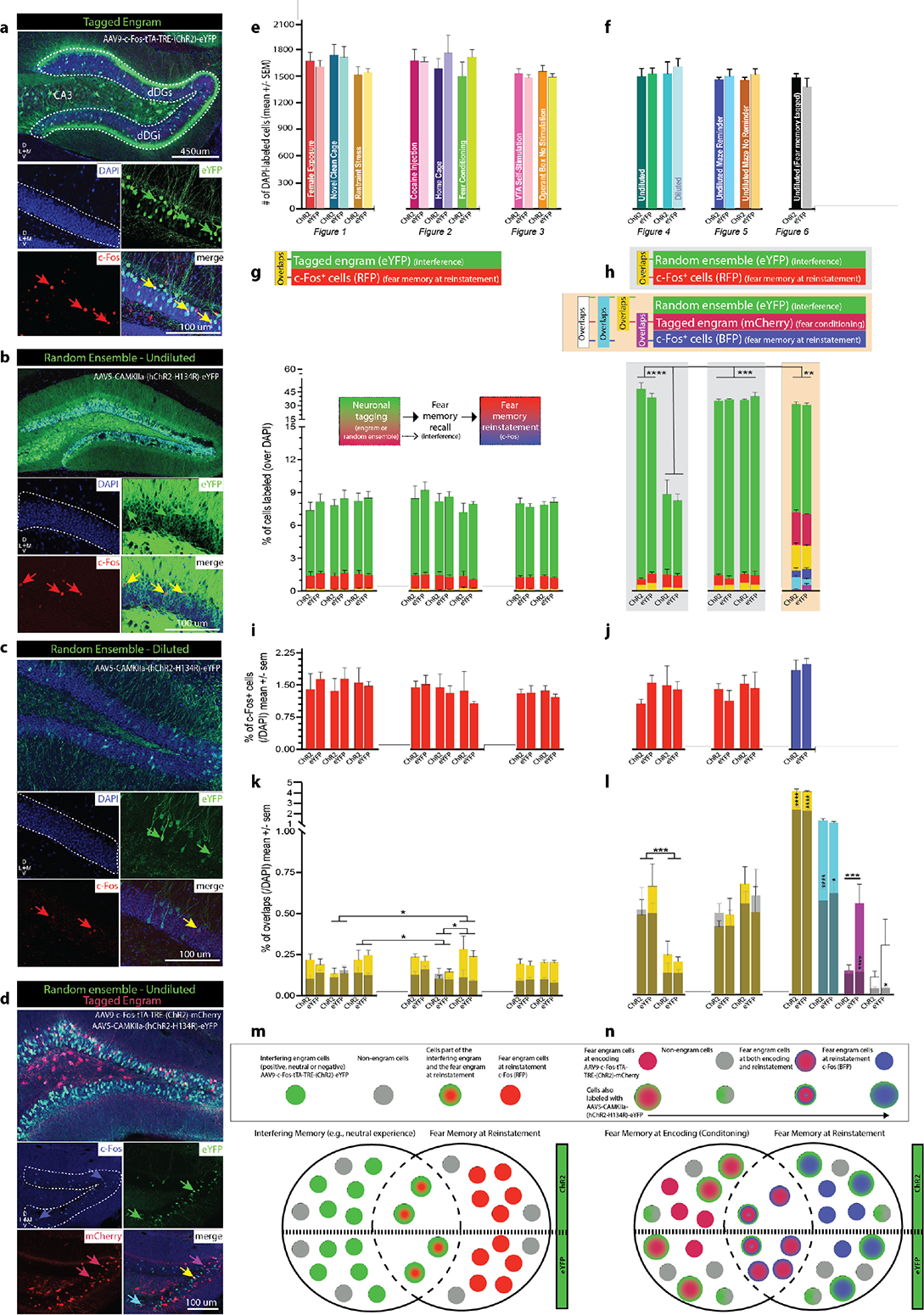
Perturbation of dDG neurons during fear memory reconsolidation rewrites the original fear memory but does not bias it in the direction of the interfering cellular ensemble. Representative images (20x) of **a,** a tagged engram in the dDG (dDGs: suprapyramidal layer; dDGi: infrapyramidal layer) using AAV9-c-Fos-tTA-TRE-(ChR2)-eYFP. DAPI (blue, enclosed within white dotted lines), eYFP (green arrows), c-Fos (red arrows), overaps (yellow arrows). **b,** Randomly labeled neurons in the dDG injected with undiluted AAV5- CaMKIIa-(hChR2-H134R)-eYFP. DAPI (blue), eYFP (green), c-Fos (red), overaps (yellow). **c,** Randomly labeled neurons in the dDG injected with diluted AAV5-CaMKIIa-(hChR2-H134R)- eYFP. DAPI (blue), eYFP (green), c-Fos (red), overaps (yellow). **d,** Both, a tagged engram in the dDG using AAV9-c-Fos-tTA-TRE-(ChR2)-mCherry and randomly labeled neurons in the dDG injected with undiluted AAV5-CAMKIIa-(hChR2-H134R)-eYFP. mCherry (red), eYFP (green), c-Fos (blue), mCherry-eYFP overaps (yellow), mCherry-c-Fos overlaps (magenta), eYFP-c-Fos overlaps (cyan). **e,** Number of DAPI-labeled neurons in each group across experiments where we tagged heterogeneously-valenced engrams to interfere with fear reconsolidation. **f,** Number of DAPI-labeled neurons in each group across experiments where we tagged randomly labeled ensembles to interfere with fear reconsolidation. **g,** The percentage of cells (/DAPI) labeled with eYFP (green, tagged engram), RFP (c-Fos^+^, red, fear memory at reinstatement), or both (overlaps, yellow) corresponding to the graph above. **h,** Left & Middle: The percentage of cells (/DAPI) labeled with eYFP (green, random ensemble), RFP (c-Fos^+^, red, fear memory at reinstatement), or both (overlaps, yellow) corresponding to the graph above. The percentage of cells in the dDG labeled with diluted virus was similar to the size of a tagged engram and significantly lower than dDG cells labeled with undiluted virus (two-way ANOVA: F(4,54)=27.55, P<0.0001, Group). Right: The percentage of cells (/DAPI) labeled with eYFP (green, random ensemble), mCherry (tagged FC engram), or both (overlaps, yellow), BFP (c-Fos^+^, blue, fear memory at reinstatement), eYFP and BFP (overlaps, cyan), and mCherry and BFP (overlaps, magenta) corresponding to the graphs above. **i-j,** The percentage of c-Fos^+^ cells (/DAPI) labeled with RFP or BFP enlarged. **k,** The percentage of overlaps (/DAPI) corresponding to the graphs above, enlarged and set against chance (grey). There was a significantly higher degree of overlap between the fear memory at reinstatement and engrams associated with negative experiences compared to engrams associated with neutral experiences despite viral condition (Chr2 or eYFP) (restraint stress vs. homecage: P=0.0461, FC vs. homecage: P=0.0121, FC vs. novel clean cage: P=0.0018) (two-way ANOVA: F(7,131)=3.427, P=0.0021, Group). Overlaps were not significantly higher than chance suggesting that our manipulation did not bias the fear engram in the direction of the interfering engram even when behavioral fear was significantly reduced. **l,** The percentage of overlaps (/DAPI) corresponding to the graphs above, enlarged and set against chance (grey). Left & Middle: Overlaps were higher when mice were injected with undiluted virus compared to diluted virus (three-way RM ANOVA: F(1,26)=7.356, P=0.0117, Dilution x Chance), a direct result of more cells being labeled. However, they were not significantly higher than chance suggesting that the cells we randomly labeled did not disproportionately include cells involved in the fear engram at reinstatement meaning those cells did not become the ensuing fear engram. Right: The percentage of cells that were part of the original fear memory and also part of the fear memory at reinstatement are significantly lower in ChR2-mice (magenta). This suggests that our reconsolidation-interference manipulation caused a disengagement of these ensembles in an orthogonal manner, separating these memories to the degree that you would see overlap in two differentially-valenced memories (P=0.0005) or simply chance levels (P<0.0001) (two-way RM ANOVA: F(1,11)=15.70, P=0.0022, Virus x Chance). The proportion of cells we randomly labeled / activated (green) that were also part of the original fear engram (cherry) was also above chance (yellow, ChR2: P<0.0001, eYFP: P<0.0001) (two-way RM ANOVA: F(1,11)=155.3, P<0.0001, Chance). However, the proportion of cells we randomly labeled / activated (green) that were also part of the fear engram at reinstatement (blue) was also above chance (cyan, ChR2: P<0.0001, eYFP: P=0.0103) (two-way RM ANOVA: F(1,11)=48.46, P<0.0001, Chance). **m-n,** Schematic to depict the ensemble dynamics described in k-l. All data are represented as means ± s.e.m. *P<0.05, **P<0.01, ***P<0.005, ****P<0.00. dDG: dorsal dentate gyrus, FC: fear conditioned.

## Discussion

Here, we combined our viral neuronal tagging strategy with optogenetics to manipulate hippocampal ensembles and disrupt the expression of a fear memory in mice. We showed that hippocampal interference induced by optical reactivation of a competing, positive memory was sufficient to disrupt reconsolidation of a fear memory. While it is generally considered evolutionarily advantageous to remember emotionally significant events well, the pursuit of therapeutic forgetting has emerged in cases such as PTSD where these memories become debilitatingly intrusive. Currently, the majority of pharmacological and cognitive behavioral treatments used to treat disorders of emotional memory typically only affect the strength of the affective response while the original fear memory is left intact^48^. As a result, these memories often recover their strength following subsequent aversive events involving stress. Studies testing reconsolidation theory have provided experimental evidence that memories are not immutable and can be updated if certain conditions are met^4, 49^. One condition is that the memory must be reactivated and secondly, that the treatment aimed at altering the memory must happen post-reactivation^50^. Our reconsolidation-interference manipulation, aimed at updating the original fear engram with attributes of a positively valenced, tagged memory, allowed for both these requirements to be satisfied. However, to limit the optical stimulation period, we designed our first experiment to test whether the stimulation protocol would be more effective when administered in the first or last half of the session. We were surprised to find our manipulation worked more effectively when it occurred simultaneously with memory recall as opposed to ten minutes post-reactivation. We speculate that the efficacy of our stimulation may be disrupted when it occurs later in the session as a result of our manipulation acting on extinction-like processes as the strength of the conditioned fear decreases across the session.

Artificially reactivating a previously consolidated memory likely leads to reconsolidation of that trace as well as that of the naturally recalled fear memory. While we did not test this directly, it is this process involving plasticity that potentially confers the activated memory with the capacity to interfere with and modify the expression of the fear memory. We found this strategy was more effective at reducing fear when the competing engram was associated with a positive experience. These effects were observed in real-time, and they enhanced rates of extinction learning, prevented stress-induced reinstatement, and persisted two weeks later demonstrating the enduring nature of our manipulation. Contrastingly, reactivation of an engram associated with a negative or neutral experience, with the exception of exposure to a novel clean cage, was not sufficient to diminish freezing when assessed immediately after stimulation, upon stress-induced reinstatement, or during spontaneous recovery (Fig. 8). These results, which were not related to differences in perturbed population sizes^30^ or locomotion, corroborate our previous findings^27^ and highlight the importance of valence. Of note, it is possible the engrams associated with negative experiences, especially fear conditioning, are highly similar to the fear memory thereby providing less interference. Indeed, we observed that dDG cells processing negative memories overlapped more with the fear memory at reinstatement in comparison to the neutral memories. Although our effects persisted two weeks later, one potential limitation of this study is that we did not attempt to reinstate a fear memory using immediate shock at a remote time point as this is similar to stress-induced reactivation of pathological conditioned fear in PTSD, which future studies may further delineate.

**Fig. 8.**
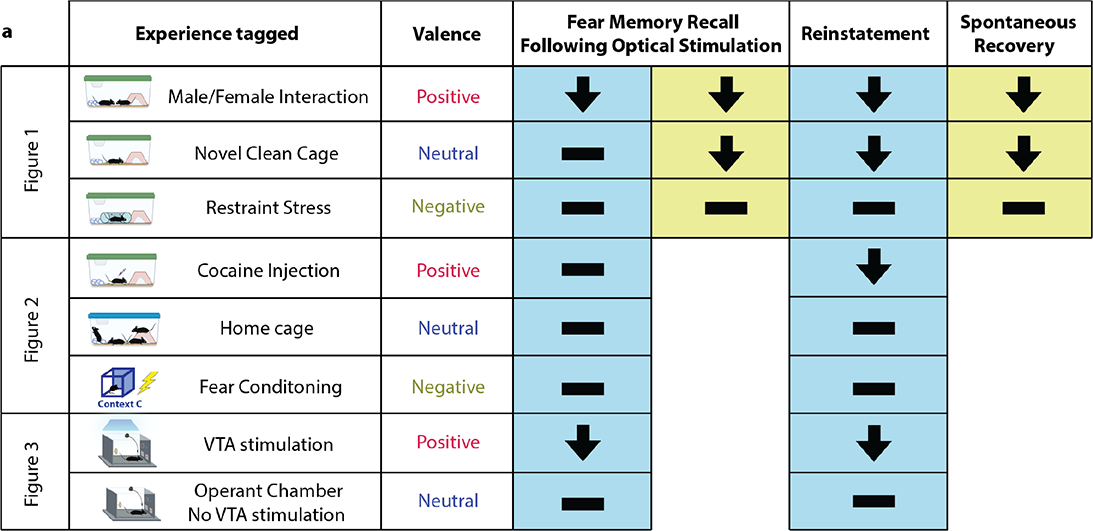
The significance of valence. **a,** Reactivation of a competing positive memory via optical stimulation is sufficient to interfere with reconsolidation of a fear memory. Reactivation of an engram associated with a negative experience, or a neutral experience with the exception of exposure to a novel clean cage, was not able to diminish freezing levels when assessed immediately after stimulation, upon stress-induced reinstatement, or spontaneous recovery.

We included closed-loop self-stimulation of the VTA as a positive experience to obtain a quantifiable measure of positive valence associated with reward, and to probe motivational aspects of operant responding for reactivation of a positive memory. Our results revealed the inherently reinforcing nature of experiencing positive affect even when it is artificially induced. Interestingly, we also observed a suppression of fear in dDG-eYFP control mice that underwent VTA self-stimulation. While these mice did not have this experience reactivated, they still experienced a reduction in fear. In humans, there is evidence to show that trait positive affect can protect against stress influencing health outcomes^51^, an effect known as the undoing hypothesis^29^. In line with this hypothesis, these results suggest hippocampal involvement in processing emotional memory may contribute to its regulation of stress responses.

Activating a randomly labeled set of neurons in the dDG was sufficient to disrupt fear memory reconsolidation. Moreover, activating 40% rather than 8% of cells yielded the most robust effects on freezing. We believe our manipulation works similarly to other stimulation-based interference protocols associated with neural plasticity such as electroconvulsive therapy (ECT), deep-brain stimulation, or trans-magnetic stimulation. While there is a paucity of literature on these treatments in the context of conditioned fear with mixed effects reported^52^, it is possible the timing of the manipulation is a key factor for treatment efficacy. For instance, stimulation occurring concurrently with recall of the fear memory may provide a unique window of opportunity to leverage reconsolidation mechanisms. Importantly, while one major side effect of ECT is the nondiscriminatory manner in which amnesia is induced^53^, our approach yielded a highly specific effect on the fear memory itself, as our manipulation did not affect a separate spatial memory, suggesting our reconsolidation-interference procedure is specific to the memory recalled at the time of stimulation.

Mechanistically, we found no evidence that the competing engram or interfering ensemble was merged with the fear memory. That is to say, the cells active during reinstatement did not contain a significant proportion of cells that were part of the ensemble we artificially activated. Only in the experiment where we determined the original fear memory was altered did we observe a high degree of overlap with both the original fear memory and the interfering ensemble. Overlap between the fear memory at reinstatement and the original fear engram was significantly reduced in experimental animals suggesting a causal relationship between reconsolidation-interference and the disengagement of these ensembles, perhaps reflecting a cellular correlate of permanently altering the fear memory.

Overall, our findings point to the dDG as a potential therapeutic node with respect to artificially modulating memories to suppress fear. In future experiments, we will examine the usefulness of these manipulations in other brain regions implicated in PTSD (e.g., amygdala) and explore the development and refinement of novel modulation strategies. Alleviating cellular, circuit-level, and behavioral abnormalities comprising memory updating impairments and maladaptive conditioned behavioral states involved in disorders such as PTSD, we believe, has promising clinical significance.

## Methods

### Animals

All experimental procedures were conducted in accordance with protocol 2018000579 (17-008) approved by the Institutional Animal Care and Use Committee at Boston University. Experimental mice included 256 wild-type (WT) male c57BL/6 mice (∼39 days of age; Charles River Labs) weighing 20–22 g at the time of arrival. Additionally, 26 WT female c57BL/6 mice (∼39 days of age; Charles River Labs) weighing 18–20 g at the time of arrival were used for female exposure (Fig. 1). Two DAT ^IRES-cre^ knock-in breeding pairs of mice were purchased from The Jackson Laboratory (B6.SJL-Slc6a3tm1.1(cre)Bkmn/J Stock No. 006660) and used to maintain an in-house breeding colony. For genotyping, a mouse tail biopsy was performed in accordance with the Boston University IACUC and 201800579 (17-008) protocol. From this colony, 50 male transgenic mice were used for the study. Mice were housed in groups of 2–5 per cage. All mice were kept on a regular light cycle 12:12 h light–dark in a temperature and humidity-controlled colony room. Cages were changed once a week and contained cardboard huts and nesting material for enrichment. Upon arrival in the facility, all mice were placed on a 40 mg/kg DOX diet (Bio-Serv, product F4159, Lot 226766) and left undisturbed for a minimum of 3 d prior to surgery with *ad libitum* access to food and water.

### Stereotaxic surgery

Aseptic surgeries were carried out with mice mounted in a stereotaxic frame (Kopf Instruments) with the skull flat resting on a heating pad. They were anesthetized with 4% isoflurane and 70% oxygen (induction), and isoflurane was reduced to 2% thereafter (maintenance). All viral constructs and coordinates are discussed below. Viral infusions were administered via a 10 μL gas-tight Hamilton syringe attached to a micro-infusion pump (UMP3, World Precision Instruments) which occurred at a rate of 100 nL min^−1^. The infusion needle was left in place for 2 min following each infusion to avoid liquid backflow. Following injections, optic fibers were implanted and secured with two anchor screws, and a mixture of metabond and dental cement to build a head cap. All mice received 0.2 mL physiological sterile saline (0.9%, s.c.), 0.1 mL of a 0.03 mg/mL buprenorphine solution (i.p.), and meloxicam (5mg/kg, s.c.) at the beginning of surgery. At the end of surgery, mice were placed on a heating pad and given hydrogel in addition to *ad libitum* food and water. Mice were given an additional injection of buprenorphine (0.1 mL, 0.03 mg/mL, i.p.) 8-12 hours later, and 8-12 hours after that both meloxicam (5mg/kg, s.c.) and buprenorphine (0.1 mL, 0.03 mg/mL, i.p.). With the exception of cage changes and being weighed, wildtype mice were left undisturbed for a 10 d period following surgery to allow for recovery and virus expression. For transgenic mice, viral expression took 6-8 weeks.

### Viral Microinjections

For experiments where we labeled the cells involved in a behavioral epoch [i.e., appetitive (positive), neutral, aversive (negative)] (Fig. 1-2, 6), mice received bilateral infusions of a viral cocktail of pAAV9-cFos-tTa (UMass Vector Core - titre: 1.5×10^13 GC/mL) and pAAV9-TRE- (ChR2)-eYFP (UMass Vector Core - ChR2 & eYFP titre: 1×10^13 GC/mL) or pAAV9-TRE- (ChR2)-mCherry (UMass Vector Core - ChR2 titre: 1.1×10^13 GC/mL, mCherry titre: 1×10^13 GC/mL) in a volume of 300 nL/side at AP: -2.2, ML: ±1.3, DV: −2.0 (relative to Bregma in mm) into the dorsal dentate gyrus (dDG) and bilateral optic fibers were implanted (AP: -2.2, ML: ±1.3, DV: −1.6, relative to Bregma). For experiments where we randomly activated cells in the dDG, mice were infused with diluted (1:5000) or undiluted virus pAAV5-CAMKIIa-(hChR2-H134R)- eYFP (Addgene-ChR2 titre: 1×10^13 GC/mL; UNC Vector Core - eYFP titre: 3.6×10^12 GC/mL) (300 nL/side) bilaterally into the same coordinates as above (Fig 4-6).

For experiments where DAT-Cre mice delivered closed loop stimulation into the left ventral tegmental area (VTA) (Fig 3), (AP: -3.1, ML: -0.5, DV: −4.4) we infused 450nL of pAAV5-Ef1a- DIO-(hChR2-E123A)-eYFP (Addgene-ChR2 & eYFP titre: 2.2×10^13 GC/mL)^54^. For these mice, bilateral infusions of the AAV9-cFos-tTa-(ChR2)-eYFP virus were delivered into the dDG. Due to space constraints, these were delivered at AP: -1.9, ML: ±1.18, DV: −1.8 where one optic fiber was placed over the left dDG at a 9° rostral angle (AP: -1.4, ML: -1.18, DV: −1.4), and the other optic fiber was placed straight over the right dDG (AP: -1.9, ML: +1.18, DV: −1.4). An additional optic fiber was placed straight over the left VTA (AP: -3.1, ML: -0.5, DV: −4.1).

### Experimental Design

#### Genetically Labeling dDG Neurons Involved in a Behavioral Epoch

We genetically labeled neurons that were active during distinct behavioral epochs using an activity-dependent and inducible Tet-Off (tetracycline inducible) optogenetic system^28, 30^. This system involves delivery of an adeno-associated viral (AAV) cocktail that allows for expression of a tetracycline transactivator protein as well as a tetracycline response element, that when bound allow for the expression of a light sensitive protein e.g., channelrhodopsin (ChR2) fused to a fluorescent reporter gene e.g., enhanced yellow fluorescent protein (eYFP). The system is inducible because transcription is controlled (reversibly turned on or off) by tetracycline, or more stable derivatives of tetracycline such as doxycycline (DOX)^55^ which is present in the animal’s diet. In order to label neurons involved in a particular behavioural epoch, the DOX diet is replaced with standard lab chow (*ad libitum*) 42 h prior to labeling. The system is activity-dependent because it is driven by the c-Fos promoter which has been widely used as a neuronal marker^56^. We labeled the cells active during putatively positive, neutral, and negative experiences in the dDG. Each of these experiences are described in more detail below. Following behavioral tagging, mice were returned to their home cages and again placed on a DOX diet. The next day, they were fear conditioned.

#### Fear Conditioning

Behavior was performed in conditioning chambers with cameras mounted to the roof for video recording (context A). Video was fed into a computer running Freeze Frame/View software (Coulbourn Instruments, Holliston, MA); freezing was measured and defined as a bout of immobility lasting 1.25 s or longer (Freezeview, Coulbourn). During the fear conditioning (FC) session, 4 shocks were delivered at 198 s, 280 s, 360 s, and 440 s during the 500 s session^31, 32^ (2 s duration, 1.5 mA intensity). Subjects were placed in a holding tank until all cage mates were fear conditioned before being returned to their home cage.

#### Recall

On the following day, mice were returned to the conditioning context for a 20 min recall session. In experiment 1, mice received optogenetic stimulation (10 ms pulse width, 473 nm, 20 Hz) of the genetically labeled ensemble with blue light during either the first or last 10 min of the session. In subsequent experiments, this was conducted only during the first 10 min. Optic fiber implants were attached to a patch cord connected to a blue laser diode controlled by automated software (Doric Lenses). Laser output was tested at the beginning of every experiment to ensure that at least 15 mW of power was delivered at the end of the patch cord (Doric lenses).

#### Extinction Training

Over the next two days, mice underwent 30 min extinction training sessions in the original conditioning context. For these sessions mice were not given optical stimulation or shock.

#### Immediate Shock

On the subsequent day, the animals were given an immediate shock in a new context (context B). A single shock (2 s duration, 1.5 mA intensity) was delivered in the first 2 s of the session, so that animals would not form a contextual representation of the environment and therefore not form an associative fear memory but would still experience stress^57–60^. Nevertheless, the context was distinctively different from the fear conditioning context with inserts and patterned walls, different lighting conditions, and almond extract odor present.

#### Reinstatement Test

The following day, mice were returned to the conditioning context for a 10 min immediate-shock induced reinstatement test.

#### Spontaneous Recovery

A subset of mice was not given immediate shock following extinction. These mice were left in their home cage for two weeks and then put back into the conditioning context for a 10 min recall test to assess spontaneous recovery of the conditioned freezing response.

#### Dependent Measures

For all sessions freezing levels were measured using Freezeview software (Coulbourn) except during recall when they were manually scored due to the presence of the optic cables that made automated scoring with Freezeview (Coulbourn) not possible.

### dDG-Labeled Behavioral Epochs

#### Female Exposure (Positive)

The experimental male mouse was placed into a clean cage with a cage top and bedding, which was used as the interaction chamber. A female mouse (PND 30-40) was then placed into the cage, and they were allowed to interact freely for 1 h^27, 31^.

#### Novel Clean Cage (Neutral)

Mice were placed into an empty clean home cage with bedding for 1 h^18^.

#### Restraint Stress (Negative)

Mice were placed into a restraint tube with air holes in an empty home cage with bedding for 1 h^61^.

#### Acute Cocaine Exposure (Positive)

Mice were habituated to scruffing and i.p. injections with saline during handling. Cocaine hydrochloride (Sigma) was prepared in 0.9% NaCl at a concentration of 2.5mg/ml. On the day of behavioral tagging, animals received an i.p. injection of cocaine at a dose of 15 mg/kg^35^. Immediately following the injection mice were placed into a clean cage with bedding for 1 h.

#### Home Cage (Neutral)

Mice remained undisturbed in their home cage during the tagging window.

#### Fear Conditioning (Negative)

Mice were fear conditioned using the same protocol as above, however, this was conducted in a distinct environment in a different room (context C), with a larger apparatus, different lighting conditions and cues on the chamber walls^30^.

#### Operant responding for closed loop VTA stimulation (Positive; control mice: Neutral)

Customized operant testing chambers were constructed from standard mouse fear conditioning apparatuses (Med-Associates) (context E). Custom-built nose-poke holes (opening diameter: 23 mm) were built into the right and left positions of one wall, and a plastic wheel (diameter: 60 mm) on a ball-bearing was fixed to the right position on the opposing wall. An Arduino Mega microcontroller was used to catalog nose pokes detected via infrared sensors (adafruit), and to detect wheel manipulation via a rotary encoder (US Digital). Custom Matlab code was written to deliver closed-loop stimulation, and to record nose pokes at each port (active and inactive) as well as wheel manipulations at each moment during the session. Behavioral events were recorded every 100 msec, and the turnaround time for the laser stimulation remained with the 100 msec clock period. To receive optogenetic stimulation, the mouse was required to either initiate a nose poke in the correct port or manipulate the wheel such that it turned in excess of 0.1 rev per second (or 5 revs per min) as measured continuously over a 20 ms interval. Both the nose poke and the running wheel had a lockout period of 1 sec (or 500 ms following termination of stimulation train), and the animal was required to withdraw from the nose port before receiving a subsequent stimulation. Optogenetic stimulation (10 ms pulse width, 473 nm, 40 Hz) initiated by nose poking was delivered via blue light to the VTA in 500 ms bouts. Optogenetic stimulation (10 ms pulse width, 473 nm, 20 Hz) initiated by spinning the wheel was also delivered via blue light to the dDG in 500 ms bouts. Code and data are available on Github, and per reasonable request to the corresponding author.

### Experimental Procedures

#### Habituation & Baseline

Dat-Cre mice were given two 45 min habituation sessions with no access to the nose ports, and only access to the wheel, whereby moving the wheel had no consequences. The first session was considered a habituation session for the mice to familiarize themselves with the apparatus and no dependent measures were taken. The second session was considered a baseline session where dependent measures related to the wheel were obtained. These included the number of wheel rotation bouts, the distance the wheel traveled, the number of seconds the wheel was rotated.

#### Training

Mice were given 4 d of training, one 45 min trial per day where they had access to the wheel (no consequences) and the active and inactive nose ports. Nose pokes in the active port resulted in a 500 ms (left) VTA stimulation bouts while nose pokes in the active port resulted in no stimulation. Following the 3rd training session, mice were taken off DOX and 42 h later brought back for the 4th training session. Following the end of the session, mice were placed back on DOX. During these sessions we continued to obtain dependent measures related to the wheel and also measured the number of nose pokes at each port as well as the number of stimulations delivered.

#### Test

The next day, mice were placed back in the operant chamber with restricted access to the nose ports and only given access to the wheel. Wheel spins during this session resulted in 500 ms dDG stimulation bouts thereby reactivating the memory of the previous training session where VTA-Chr2 mice received self-stimulation of the VTA, and VTA-eYFP mice did not. During this session we measured the number of wheel rotation bouts, the distance the wheel traveled, the number of seconds the wheel was rotated and also the number of simulations produced by spinning the wheel to assess whether mice will perform an operant task for reactivation of a positive memory. The following day all mice underwent fear conditioning (as above).

#### Genetically Labeling Random dDG Neurons Not Tied to the Encoding of a Behavioral Epoch

We genetically labeled random dDG neurons driven by the CaMKIIa promoter to allow for the expression of ChR2 fused to eYFP. For the 1:5000 dilution, we used sterile saline. These cells were then activated during the fear memory recall session. For this system, a tagging window is not required therefore mice were not taken off DOX. However, we fed them the same DOX diet for consistency.

#### Open Field

To assess whether stimulation of a cocaine engram induced locomotor activity or had any effect on anxiety-like behavior, we tagged an acute cocaine experience (15mg/kg, i.p.) off-DOX and later placed mice in an open field arena attached to a patch cord with a camera over top. The first 5 min were considered a baseline where mice received no stimulation. In the last 5 min mice received optical stimulation in the dDG (10 ms pulse width, 473 nm, 20 Hz).

#### Radial Arm Maze

To assess whether perturbing dDG cells during fear memory recall affects other types of memory, or if the manipulation is specific to the fear memory, we trained mice on a spatial reference memory task. Mice were bilaterally infused with AAV5-CAMKIIa-ChR2-eYFP in the dDG (300nl/side). Following surgery and 10 days of recovery, mice were then water-restricted so that they would be motivated to search for a water reward in the maze. We used a custom-built eight-arm radial maze made of Plexiglas (55 mm arm width, 355.6 mm arm length, center area 152.4 mm diameter) and designed a task where the goal arm stayed consistent throughout the experiment^62^. Mice were trained to search for this location by shaping their behavior over a series of trials (see procedure below). The maze was surrounded by four curtains with distinct distal visual cues to allow mice to navigate and locate the reward using these extra-maze cues.

Initially, mice were given one 5 min habituation trial where they had access to all 8 arms which were not baited, however after the trial, mice were placed in a clear plastic container and given 1ml of water in a falcon tube cap. The following day they were given 4 shaping trials (5 min each) and each time the goal arm was baited with 0.25 ml of water in a falcon tube cap at the end of the arm. The maze was designed with inserts within each arm to close off arms at their entry point and at the end goal location. On the first trial mice were given access to the goal location only (both doors in the goal arms were inserted, restricting the mice to the end of the arm). On the second trial mice were given access to the entire goal arm, but only that arm (only the door at the entry point was inserted). On the third trial mice were placed in the center of the maze and all arms were open. Finally, on the last trial animals were placed at a starting point (the end of the arm directly opposite the goal arm). During each trial the mice were allowed to drink and given extra time to look up at the cues around them. In cases where mice didn’t drink all the water within 5 min, we placed them in the clear plastic container and allowed them to drink the remaining amount. However, in most cases they drank all the water in the maze.

Training lasted between 5-14 days. Mice were given 4 trials / day. During these trials there was a webcam mounted over top of the maze so we could record and score their behavior. We were also able to see their behavior in real time on the other side of the room divided by a thick curtain. Dependent measures included: Latency to reach the goal location (s), arm deviations (number of arms away from the goal arm mice first visit), number of reference errors (number of arm entries into any arm besides the goal arm), repeated reference errors or working memory errors (re-entry into an incorrect arm). To reach our training criterion, mice had to demonstrate for two consecutive days, a latency score of under 90 s, an arm deviation score of less than 2, less than 2 reference errors and less than 2 repeated reference errors on trial 1 and on trials 2-4 calculated separately. Mice took approximately 8 days to reach this criterion. Once they did, the following day they were fear conditioned.

#### Immunohistochemistry

Mice were overdosed with sodium pentobarbital and perfused transcardially with (4°C) phosphate-buffered saline (PBS) followed by 4% paraformaldehyde (PFA) in PBS. Brains were extracted and stored overnight in PFA at 4°C and transferred to a solution of 0.01% sodium azide the next day. The solution was prepared by dissolving 5g of 10% sodium azide (Thermo Scientific) in 50 mL of 1X PBS to create a stock solution. This solution was then diluted to a 0.01% dilution by dissolving 1 mL of the 10% stock solution in 999 mL of 1X PBS.

Brains were sliced into 50 μm coronal sections with a vibratome (Leica, VT100S) and collected in cold PBS. Sections were blocked for 2 h at room temperature in 1x PBS + 2% Triton (PBS-T) and 5% normal goat serum (NGS) on a shaker. Sections were transferred to well plates containing primary antibodies made in PBS-T [1:1000 rabbit anti-c-Fos (SySy 226-003, Fig. 1-5; Abcam ab190289, Fig. 6), 1:500 rabbit anti-TH (Millipore AB152, Fig. 3), 1:1000 chicken anti-GFP (Invitrogen a10262, Fig. 1-6), or 1:1000 guinea anti-RFP (SySy 390 004 Fig. 6)] and incubated on a shaker at 4°C for 48 h. Sections were then washed 3x (10 min) in PBS-T followed by a 2 h incubation with secondary antibodies made in PBS-T [1:200 Alexa 555 anti-rabbit (Invitrogen A21428, Fig. 1-5), 1:200 Alexa 488 anti-chicken (Invitrogen, A11039, Fig. 1-6), 1:200 Alexa 555 anti-guinea (Invitrogen, A21435, Fig. 6), 1:200 Alexa 405 anti-rabbit (Abcam ab175653, Fig. 6)]. Following 3 additional 10 min washes in PBS-T, sections were mounted onto micro slides (VWR International, LCC). Nuclei were counterstained with DAPI added to Vectashield HardSet mounting medium (Vector Laboratories, Inc). Slides were then coverslipped and put in the fridge overnight to cure. The following day the edges were sealed with clear nail-polish and the slides were stored in a slide box in the fridge until imaging.

#### Fluorescent Confocal Image Acquisition and Quantification

Images were collected from coronal sections using a fluorescent confocal microscope (Zeiss LSM 800 with airyscan) at 20x magnification. For quantification of overlaps for animals receiving bilateral viral dDG injections, 3 z-stacks (step size 0.94 μm) were taken per hemisphere from 3 different slices yielding ∼6 total z-stacks per animal. Data from each hemisphere was then pooled and the means for the 6 z-stacks were computed. These means were then used to obtain a group mean. For histological verification of VTA injections and to confirm that the VTA neurons we labeled were indeed dopaminergic, we assessed the degree to which eYFP^+^ cells were colocalized with TH^+^ cells. Mice receiving unilateral viral injections in the VTA had 3 z-stacks (step size 0.94 μm) obtained from 3 different slices yielding 3 stacks per animal. For all images, the total number of DAPI positive (+), and eYFP^+^ neurons were counted using Image J/ Fiji software (https://imagej.nih.gov/ij/). c-Fos^+^ neurons were stained with either RFP (Fig. 1-5) or BFP (Fig. 6) and quantified. mCherry+ neurons were also quantified (Fig. 6). Percentage of immunoreactive (eYFP, mCherry, RFP or BFP) neurons, including overlaps, was defined as a proportion of total DAPI-labeled cells. Chance overlap was calculated as the percentage of the first immunoreactive neuron (e.g., eYFP^+^) multiplied by the percentage of the second immunoreactive neuron (e.g., c-Fos^+^) over the total number of DAPI neurons.

#### Statistical Analyses

Calculated statistics are presented as means ± standard error of the mean (s.e.m.). To analyze differences, we used one, two and three-way analysis of variance (ANOVA) and in the cases where these are repeated measures (RM) analyses, it is stated. In some cases, we used paired t-tests. When appropriate, follow-up post hoc comparisons (Tukey’s HSD) were conducted. All statistical tests assumed an alpha level of 0.05. For all figures, * = P < 0.05, ** = P < 0.01, *** = P < 0.001, **** = P<0.0001.

## Acknowledgements

We thank Lauren Reynolds, Emily Doucette, Daniel Sheehan, Moriah White, Heloise Leblanc, Amy Monasterio, and Ryan Senne for technical assistance. This work was supported by an NIH Early Independence Award (DP5 OD023106-01), an NIH Transformative R01 Award, a Young Investigator Grant from the Brain and Behavior Research Foundation, a Ludwig Family Foundation grant, the McKnight Foundation Memory and Cognitive Disorders Award, and the Center for Systems Neuroscience and Neurophotonics Center at Boston University.

## Author Contributions

S.L.G. J.H.B., and S.R. designed the experiments. S.L.G. and A.H.F. collected the data for all experiments / figures. Data collection for: Fig. 1 was assisted by Y.Z.; Fig. 2 by E.R. & C.C.; Fig. 3 by J.H.B., L.F.R., & M.S.; Fig. 4 by C.C., E.R., & L.F.R.; Fig. 5 - 6 by A.G..; Fig 7 by L.F.R. & A.G. Data analysis and figures were completed by S.L.G.. The manuscript was written by S.L.G. and edited by J.H.B., E.R., A.H.F. and S.R.. Final edits were made by all authors.

---

**Supplementary Fig. 1.**
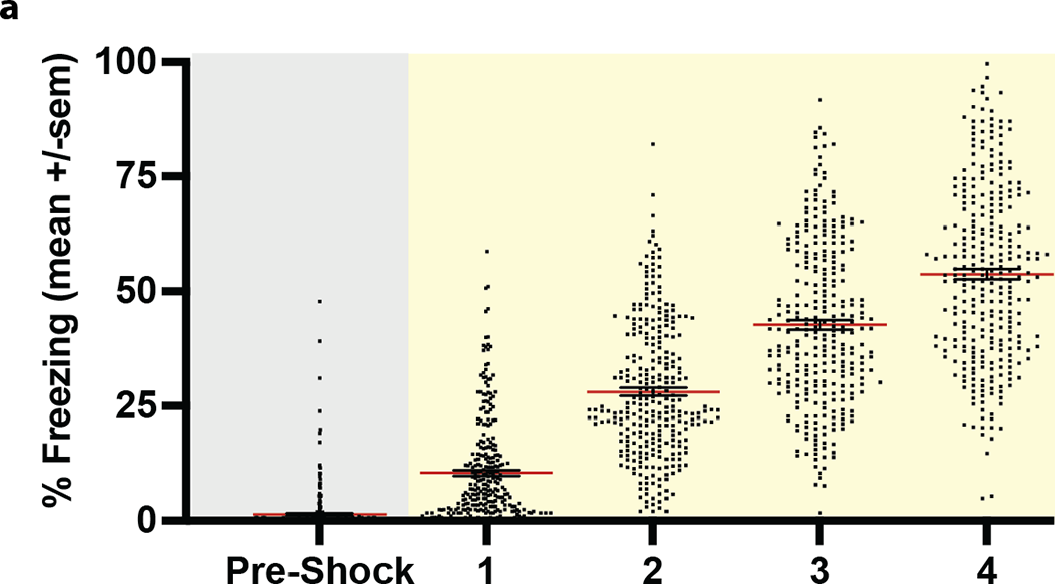
A stepwise function of fear. **a,** Mice were subject to a 4-shock fear conditioning protocol (n=297, all experiments combined). They demonstrated relatively little freezing prior to the first shock, and then gradually increased freezing levels with the presentation of each successive shock (one-way RM ANOVA: F(4,1480)=922.17, P<0.0001, Time). All data are represented as means ± s.e.m. *P<0.05, **P<0.01, ***P<0.005, ****P<0.00.

**Supplementary Fig. 2.**
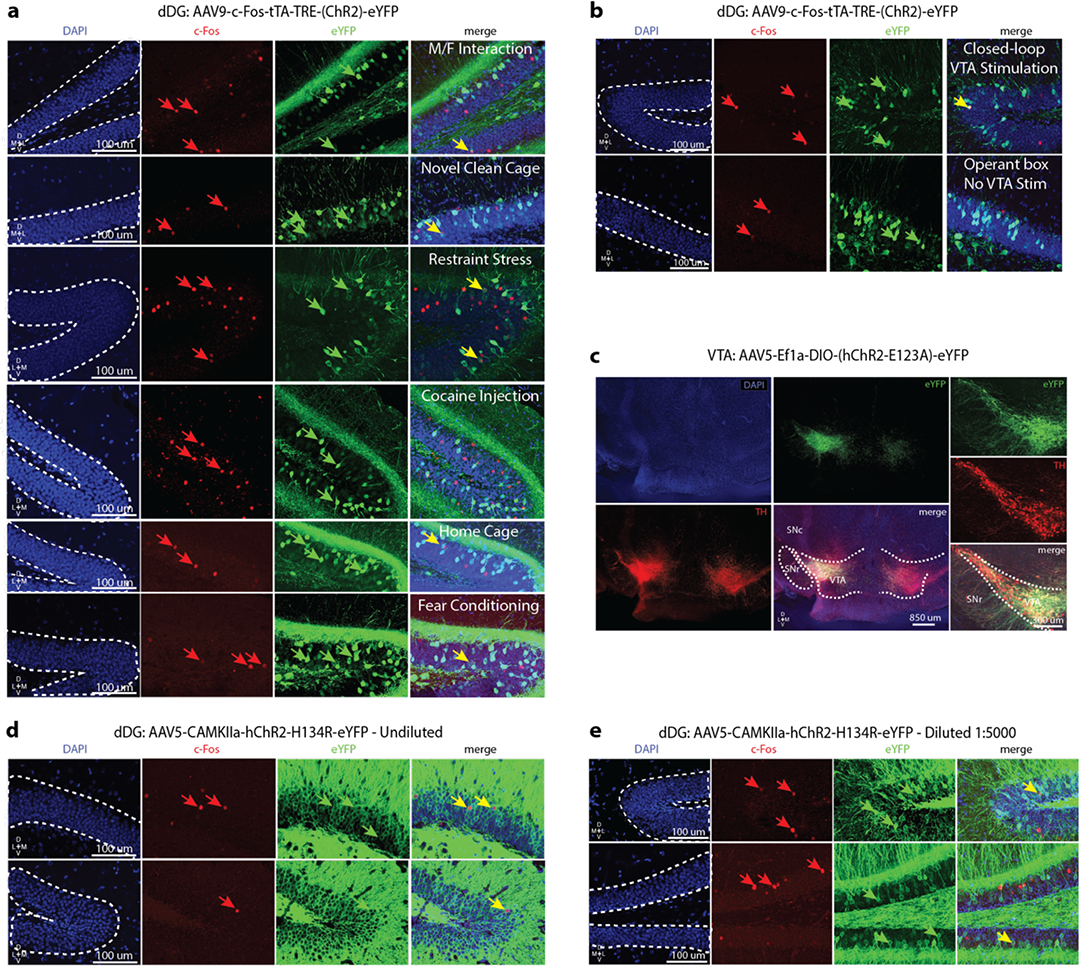
Representative images from each group. **a-b,** Representative images for dDG cells involved in the encoding of heterogeneously-valenced behavioral epochs (positive, neutral, and negative experiences labeled with AAV9-c-Fos-tTA-TRE-eYFP. Counterstain DAPI (blue), eYFP (green arrows), c-Fos (red arrows), overlaps (yellow arrows). **c,** Representative images for VTA neurons (left hemisphere) labeled with AAV5-Ef1a-DIO- (hChR2-H134R)-eYFP (green) and co-localized with tyrosine hydroxylase (TH, red), overlaps (yellow), counterstain DAPI (blue). **d-e,** Representative images for dDG cells randomly labeled with undiluted and diluted AAV5-CaMKIIa-(hChR2-H134R)-eYFP. Counterstain DAPI (blue), eYFP (green arrows), c-Fos (red arrows), overlaps (yellow arrows). dDG: dorsal dentate gyrus, SNc: substantia nigra pars compacta, SNr: substantia nigra pars reticulata, TH: tyrosine hydroxylase, VTA: ventral tegmental area.

